# Saddle Curvature Association of nsP1 Facilitates the Replication Complex Assembly of Chikungunya Virus in Cells

**DOI:** 10.1101/2025.05.01.641776

**Authors:** Xinwen Miao, Michelle Cheok Yien Law, Jatin Kumar, Choon-Peng Chng, Yongpeng Zeng, Yaw Bia Tan, Jiawei Wu, Xiangfu Guo, Lizhen Huang, Yinyin Zhuang, Weibo Gao, Changjin Huang, Dahai Luo, Wenting Zhao

## Abstract

The replication of positive-sense RNA viruses, including the pathogenic SARS-CoV-1 and -2, DENV, and ZIKV, often takes place in curved cellular membrane compartments of tens to hundreds of nanometers in diameter within host cells. The non-structural proteins (nsPs) of viruses are found to critically control the formation and maintenance of such unique nanoscale membrane structures. However, the molecular mechanism underlying the association and assembly of nsPs around curved membranes has remained elusive, due to the technical difficulties of imaging the nanoscale interaction in live cells. In this study, we leveraged vertically aligned nanostructures to define membrane curvatures in live cells and investigated the assembly of viral nsPs on these curved membranes. Using Chikungunya virus as a model system, we found that its nsP1 was preferentially bound to and stabilized around positively curved membrane sites and the preference became more apparent as the radius of curvature decreased to 150 nm or smaller. This preferential accumulation was mainly attributed to the hydrophobic residues on recently identified membrane association loops (MA loops) of each nsP1 monomer. Molecular dynamics simulations further confirmed the improved binding kinetics and stability of nsP1 on positively curved membranes, especially when a 12-mer ring was formed by the nsP1 dodecamer. More interestingly, a saddle curvature association was enabled by the 3D coordination of nsP1 monomers along the dodecamer ring, where the MA loops of individual nsP1 monomers sensed a wide range of the positively curved membrane in the x-z plane while the rigid nsP1 dodecamer ring stabilized the negative curvature in the x-y plane. Its strong coupling to the saddle curvature fulfilled the need to constrain the neck of the membrane spherule for viral replication. Strikingly, productive CHIKV replication exhibited strong enrichment on patterned nanostructure array, proving the effectiveness of membrane curvature-guided assembly of the functional replication complex. Our findings revealed that the cell membrane facilitated the local enrichment of viral nsPs in a curvature-dependent manner. It opens up membrane curvature modulation as a new dimension to dissect the formation and regulation of membrane compartments for viral replication.

## Introduction

Many positive-strand RNA viruses generate nanoscale confined membrane compartments dedicated to protecting and facilitating their genome replication. These include the highly pathogenic viruses like hepatitis C virus (HCV),^1^ dengue virus (DENV),^2^ zika virus (ZIKV),^3^ the severe acute respiratory syndrome coronavirus (SARS-CoV),^4^ and the latest SARS-CoV-2^5^ causing the COVID-19 pandemic. Such viral replication-associated membrane compartments have been found in various structures composed of either double- or single-membrane vesicles, ranging from 40 nm to hundreds nm,^6^ and extending from diverse cellular membranes including both plasma membrane (PM) at the cell surfaces^7^ and the intracellular organelles like endoplasmic reticulum (ER),^8^ mitochondria,^9^ endosomes,^10^ lysosomes,^11^ and peroxisome.^12^ However, how viruses hijack the host cell membrane to assemble specific nanoscale membrane compartments for their own purpose is largely unclear. These membrane compartments are usually associated with viral replication complexes (RCs) formed by non- structural proteins (nsPs), together with unknown host factors in many cases, to help RNA synthesis.^13,14^ Among them, only a handful of nsPs are responsible for the binding and remodeling of the host cell membrane.^15^ They act on a variety of membrane shapes, i.e. from highly curved ER tubular networks^16^ to spherule-shaped mitochondria^9^ and PM.^7^ Geometrically, nsPs are also found on multiple curvature types including not only the positive and negative curvature in a 2D plane^17,18^ but also the unique saddle curvature in 3D.^13,14^ One intriguing yet unexplored question is whether or how the existing curvatures predispose or modulate the recruitment and organization of nsPs for membrane remodeling during viral replication.

Various technologies have been explored to study the nanoscale assembly of viral nsPs on host cell membranes. Electron microscopy (EM) is the most popular method to visualize host cell membranes remodeled by viral nsPs with nanometer resolution.^2,8,10^ However, the location of viral nsPs is poorly resolved with immunogold labeling and no dynamic correlation between protein and membrane curvature is achievable. Transient expression of fluorescent-tagged viral nsPs in host cells has been widely used to examine their dynamic distribution on^19,20^ or co-localization with various intracellular organelles.^21,22^ Unfortunately, the formation of nanoscale membrane compartments is often below the optical resolution of fluorescence microscopy and thus is hardly resolved beyond diffraction- limited puncta. Liposome binding and co-floating assays are used to measure nsPs’ binding affinity to the curved vesicles *in vitro*,^23,24^ where the binding-related protein conformational changes can also be measured by circular dichroism (CD) spectroscopy,^25^ and the resulting membrane deformation if any is visualized by TEM.^25–27^ However, whether the *in vitro* behaviors reflect the native viral replication complex (RC) assembled in live cells needs to be validated.

Recently, direct manipulation of nanoscale membrane curvature in cells has been achieved using nanostructure-based surface engineering assay^28–35^ and optical tweezer-based membrane tethering assays.^36–38^ These approaches have demonstrated that nanoscale membrane curvature can enhance the accumulation of endocytic proteins,^28,32^ Ras clustering,^30^ G protein-coupled receptors,^30,36^ glycoproteins,^33^ Bin/Amphiphysin/Rvs (BAR) proteins,^39^ and Wiskott–Aldrich syndrome protein (WASP)^35^, as well as facilitate the actin polymerization^29^ and the integrin-mediated adhesions^34^. Here, we fabricated vertically aligned nanostructure arrays with various designed geometries to provide both positively and negatively curved PM sites with defined nanoscale curvature values in live cells. This platform allows for the survey of a range of curvature values and signs that are likely presented at different stages during the progressive formation of highly curved membrane spherules from relatively flat membranes. Taking Chikungunya virus (CHIKV) as a model, we probed the distribution of its membrane-targeting nsP1 on both positively and negatively curved membranes. The impact of lipid composition and different protein domains were examined in live cells and validated *in vitro* using supported lipid bilayers (SLBs) formed on the nanostructures. All-atom and coarse-grained simulations were performed to evaluate the contribution of curvature to the binding and stabilization of nsP1 on the membrane. Furthermore, membrane curvature-guided viral replication complex assembly in infected cells was successfully demonstrated using patterned nanostructure arrays.

## Result and Discussion

### 1. Positive curvature-promoted membrane association of CHIKV nsP1 in live cells

Different from other alphaviruses, the membrane spherules for CHIKV replication are mainly observed on PM instead of cytopathic vacuoles.^40,41,7^ Among the four nsPs of CHIKV that are needed for the membrane spherule formation on PM, nsP1 is the only one known to associate with the membrane, serving as the critical link to connect PM and CHIKV RC assembly.^19,42,43^ Yet, how the local PM curvature affects the membrane association of nsP1 in live cells and consequently the viral genome replication is largely unknown.

The recent cryo-electron tomography (cryo-ET) study showed that nsPs precisely assembled at the neck of the membrane spherules where a unique saddle curvature geometry (**Fig. 1a, left column**) was formed combining positive curvature (x-z plane) and negative curvature (x-y plane) in 3D.^13,14^ An intriguing question is whether the nsPs prefer binding to positive or negative curvature or both at the same time but via different planes. However, it is technically challenging to image the 3D saddle curvature at the nanoscale due to the limited microscopy resolution. To better dissect the interaction of nsP1 with this unique saddle structure, we designed two types of nanostructures to decouple 3D saddle curvature into two 2D curvatures (**Fig. 1a, middle & right columns, Fig. 1b**). Specifically, the positive curvature at the x-z plane of the saddle point was recapitulated by the positively curved nanobar ends in the x-y plane (**Fig. 1a, top row)**; while the negative curvature at the x-y plane of the saddle point was recapitulated by the negatively curved intersection of a nanocross design in the x-y plane as well (**Fig. 1a, bottom row)**. This reorientation of both positive and negative curvatures into the x-y plane allowed a better resolution to measure the curvature-guided nsP1 accumulation on PM.

**Figure 1.**
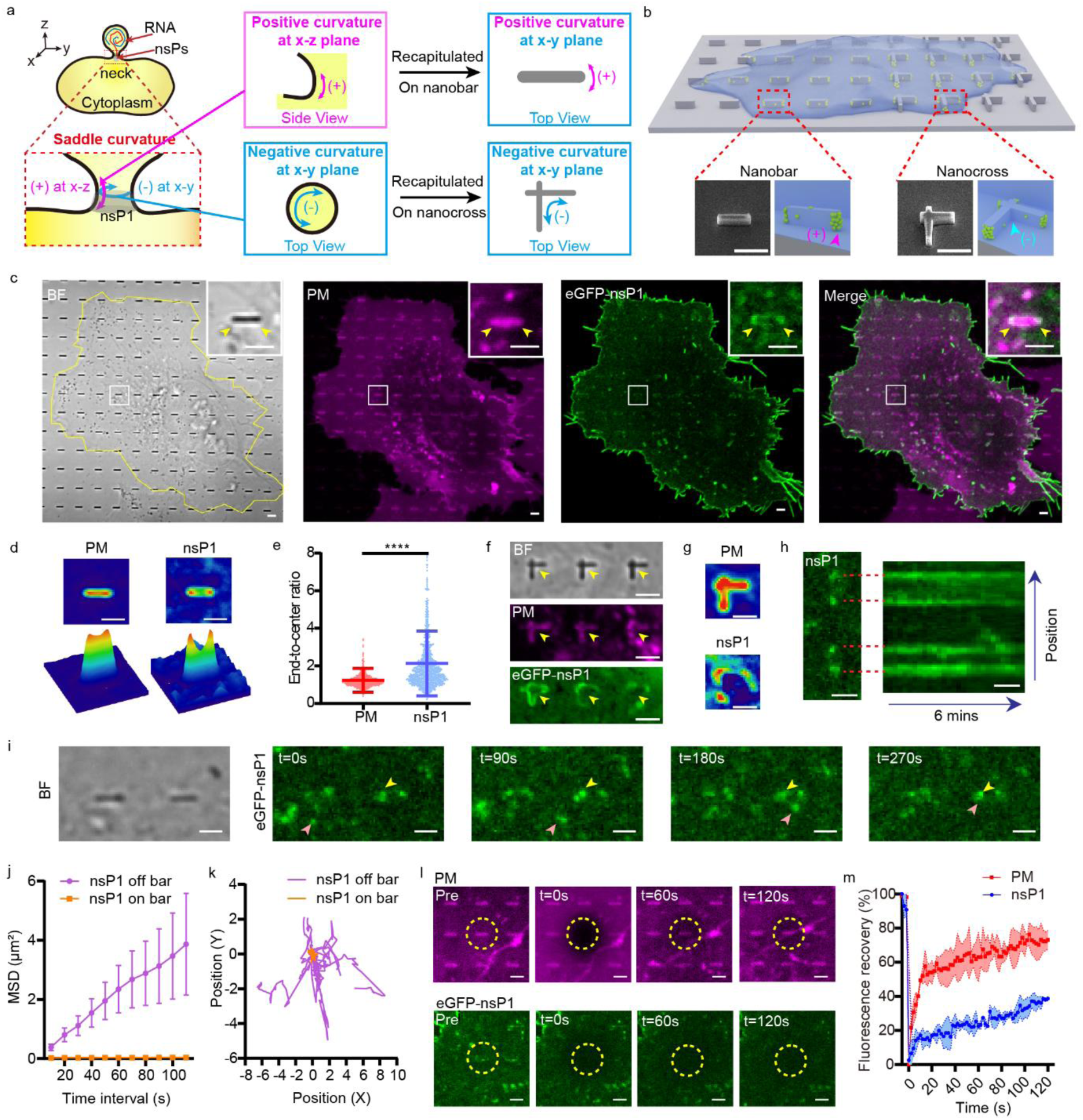
CHIKV nsP1 curvature sensing ability on nanostructures in live cells. **(a)** Schematic of a spherule on plasma membrane (PM) for CHIKV viral replication. The neck of the spherule displayed a saddle curvature with both positive (+) and negative (-) curvature but at the x-z and x-y plane correspondingly. Two types of nanostructures (nanobar and nanocross) were designed to recapitulate the 3D saddle curvature into 2D. **(b)** Illustration and the scanning electron microscopy (SEM) image of the nanostructures. The positive curvature of the nanobar was indicated by the pink arrow, while the negative curvature of the nanocross was indicated by the blue arrow. **(c)** Representative images of a U2OS cell culture on the nanobars with CellMask Deep Red stained-PM (magenta) and overexpressed eGFP-nsP1 (green), cell boundary is highlighted in brightfield. **(d)** Average images and corresponding 3D surface plots of the PM and eGFP-nsP1 distribution around nanobars. n = 489 bars for PM and n = 223 bars for eGFP-nsP1. **(e)** Scatter plot of PM and eGFP-nsP1 fluorescence intensity ratio at nanobar ends over the flat sidewall (end-to-center ratio). Data are the mean ± SD. P value calculated using unpaired t-test with Welch’s correction: **** represents p < 0.0001. n = 995 bars for PM and n = 1005 bars for eGFP-nsP1, with 3 independent experiments. **(f)** Representative images of cells cultured on the nanocross with stained PM (magenta) and overexpressed eGFP-nsP1 (green). **(g)** Average images of the PM and eGFP-nsP1 distribution around nanocross. n = 33 crosses. **(h)** Kymograph plots of eGFP- nsP1 on two adjacent nanobars. **(i)** Representative time-lapse images of eGFP-nsP1 clusters on (yellow arrows) and off (pink arrows) nanobars in 270 s. **(j)** Mean square displacement (MSD) of eGFP-nsP1 clusters on and off the nanobars in 110 s. n = 14 for clusters on the bars and n = 12 for clusters off the bars. **(k)** The trajectory of the nsP1 cluster on and off the bars in 110 s. n = 14 for clusters on the bars and n = 12 for clusters off the bars. **(l)** Representative images of confocal FRAP test for PM (magenta) and eGFP-nsP1 (green) on nanobar in 120 s. Yellow dashed circles indicate the bleaching area. **(m)** Normalized fluorescence intensity plot of the PM and eGFP-nsP1 within nanobar area from FRAP measurement (2 s interval), from 3 independent experiments. Scale bars, 2 μm.

We first examined the curvature preference of nsP1 alone using transiently expressing eGFP tagged nsP1 (eGFP-nsP1) in U2OS cells plated on nanobar and nanocross arrays (**Fig. 1b**). Interestingly, eGFP-nsP1 exhibited a significant preference to the positively curved membranes at the nanobar ends, as shown in both representative images (**Fig. 1c**) and average images (**Fig. 1d**) where the protein density at the two curved nanobar ends was significantly higher than the flat center. This curvature sensing ability of eGFP-nsP1 was mainly attributable to nsP1, rather than the fluorescence tag, as the expression of GFP only exhibited a strong cytosolic signal and lacked the ability to associate with the PM (**Supplementary Fig. S1)**. To evaluate the curvature preference of the nsP1, we quantified the fluorescence intensity at the nanobar end versus the nanobar center, indicated as ‘end-to-center ratio’ (**Fig. 1e**). A significant difference between CellMask stained membrane (1.233 ± 0.637, mean ± SD [standard deviation]) and eGFP-nsP1 (2.129 ± 1.718) was observed (unpaired t-test, p < 0.0001). The cumulative frequency distribution of the end-to-center ratio also showed a similar trend that more than 72 % of nsP1-bound nanobars displayed a ratio of more than 1 while only 36 % were observed in the CellMask labeled ones (**Supplementary Fig. S2**). Super-resolution imaging with stimulated emission depletion microscopy (STED) further confirmed that multiple nsP1 puncta were enriched at the positively curved nanobar ends (**Supplementary Fig. S3**). In comparison, eGFP-nsP1 exhibited no detectable enrichment at the negatively curved nanocross intersection, but a strong accumulation at the positively curved corners of the nanocross arms. (**Fig. 1f**, **g**). These results suggested that, although the neck of the spherule presented both positive and negative curvature sites, the assembly of nsP1 and other associated nsPs could be more dominantly affected by how curved the membrane was towards a positive direction. It is worth noting that nsP1 was also observed to accumulate inside filopodia-like protrusion structures in cells. However, recent cryo-ET studies showed that these filopodia-like structures could not generate membrane spherules for CHIKV replication,^13,14^ but were coupled with other host factors like Rac1-PAK1-Arp2/3 signaling^44^, thus less relevant to study here for the impact of curvature on spherule formation. Interestingly, we observed that CHIKV replication preferentially accumulated at the positively curved membrane of the host cells, compared to the nearby protruded area with negatively curved membranes (**Supplementary Fig. S4**), suggesting an important influence of positive curvature on CHIKV replication.

Besides the curvature-guided spatial distribution of nsP1 on the membrane, we also examined its temporal stability at curved sites. By taking a six-minute time series of eGFP-nsP1 on nanobar arrays (**Supplementary Movie 1**), we found that the nsP1 puncta were persistently associated with nanobar ends without significant intensity decay. The kymography analysis in **Fig. 1h** showed similarly that the nsP1 intensity dominated at the nanobar ends with very little signal along the flat nanobar centers, indicating that the preferential association of the nsP1 at the curved membrane sites was stable over time. In addition, the snapshots of the nsP1 puncta at the nanobar ends also showed minimal lateral movements while those on the surrounding flat membrane could still move around (**Fig. 1i**), resulting in significantly lower mean square displacement (MSD) (**Fig. 1j**) and shorter trajectory of the nsP1 puncta over time (**Fig. 1k**). The temporal analysis strongly suggested that the highly curved nanobar end could stabilize nsP1 around it. Interestingly, when we performed fluorescence recovery after photobleaching (FRAP) test on the fluorescence signals of both eGFP-nsP1 and CellMask on the nanobar ends, there was no detectable recovery observed for nsP1, but the membrane signal recovery remained normal (**Fig. 1l, m, Supplementary Movie 2**). It suggested that the nsP1 was strongly anchored at the curved PM sites with no detectable lateral diffusion and dynamic assembly.

Since the nsPs of the *alphavirus* genus are relatively conserved and share ∼60%-80% sequence homology,^40,45,46^ we further validated the curvature-guided membrane assembly of nsP1 using another virus in the same genus, Venezuelan equine encephalitis virus (VEEV). Similar to the CHIKV nsP1, VEEV nsP1 also showed a stronger membrane association on the nanobar ends than on the center (**Supplementary Fig. 5a, b)**. The end-to-center ratio of VEEV nsP1 was not significantly different compared to CHIKV (**Supplementary Fig. 5c**, CHIKV: 2.018 ± 1.871 *vs.* VEEV: 1.878 ± 1.505, unpaired t-test, p < 0.0001), with 48 % of the value higher than 1 (**Supplementary Fig. 5d**). These data indicate that the positive curvature-promoted membrane association of nsP1 is not specific for CHIKV only, but a conserved behavior among multiple alphaviruses for facilitating the viral replication process.

### 2. Recapitulating the curvature-dependent membrane association of CHIKV nsP1 *in vitro*

The positive curvature-guided membrane association of nsP1 was further validated by examining purified nsP1 on synthetic lipids bilayers formed *in vitro* on nanobar arrays as illustrated in **Fig. 2a**. Here, eGFP-fused nsP1 protein was purified from *E.coli* protein-expression system (eGFP- nsP1-(b) in short) to obtain the fraction of monomers.^45^ SLB containing 89.5% phosphatidylcholine (from egg, egg PC in short), 10% phosphatidylserine (from brain, brain PS in short) and 0.5% Texas Red™ 1,2-Dihexadecanoyl-sn-Glycero-3-Phosphoethanolamine (Texas-Red-PE) was formed on the surface of the nanobar and then incubated with eGFP-nsP1-(b). Interestingly, eGFP-nsP1-(b) monomer showed not only strong membrane binding as reported earlier,^25^ but also preferential accumulation at curved membrane sites at nanobar ends (**Fig. 2b).** The curvature preference was more obviously observed in both averaged images (**Fig. 2c**) and quantified end-to-center ratio (eGFP-nsP1: 1.109 ± 0.106 *vs.* lipid: 1.018 ± 0.121, unpaired t-test, p < 0.0001) (**Fig. 2d**). It indicated that the nsP1 sensed the membrane curvature on its own without the need for other viral materials and/or host factors.

**Figure 2.**
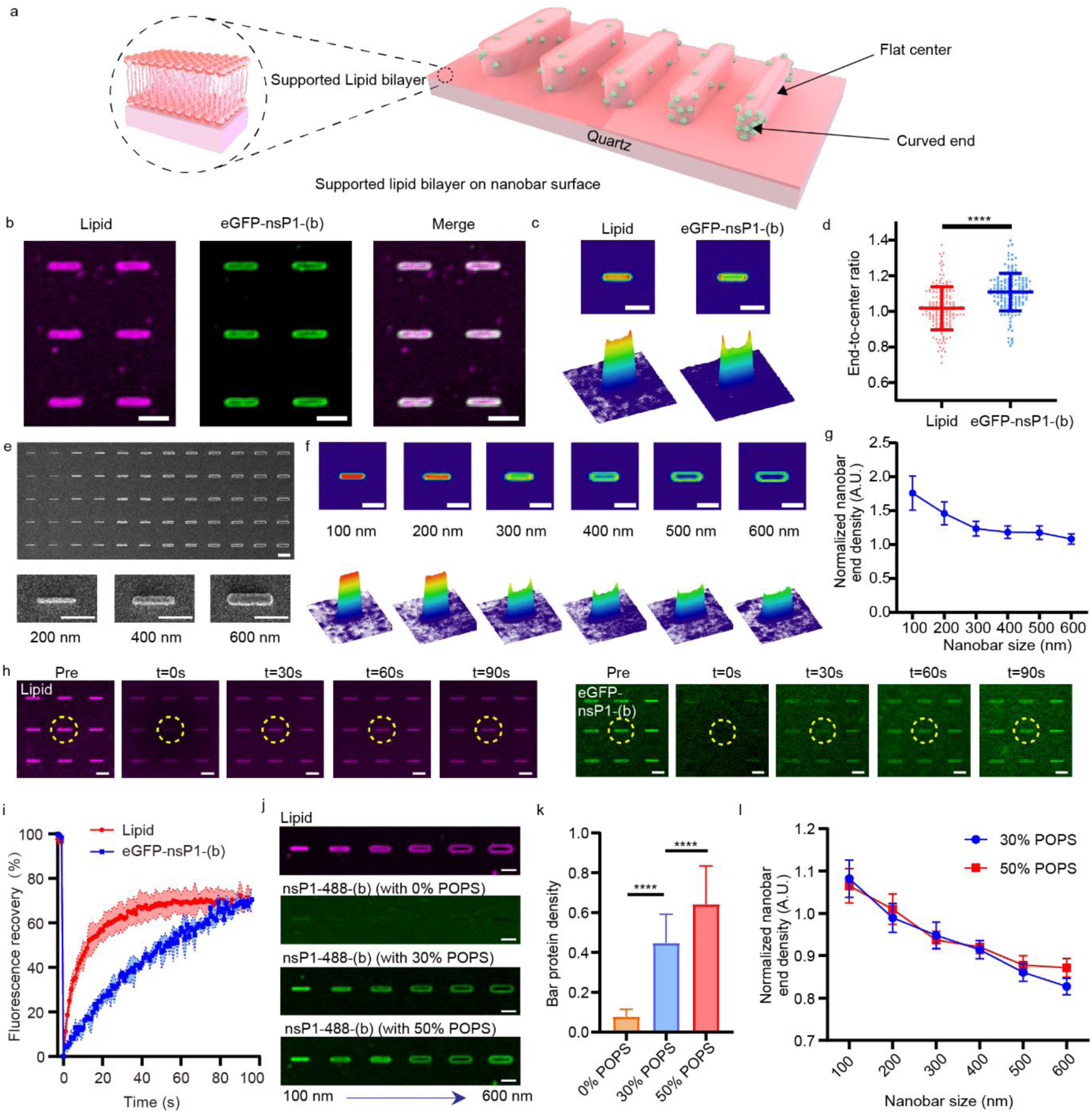
Bacterial expression CHIKV eGFP-nsP1 curvature sensing ability on supported lipid bilayer (SLB). **(a)** Illustration of the SLB wrapping around the nanobars with gradient increasing widths. Curvature sensing protein preferred to bind more around the curved ends of the smaller-sized nanobars with more curved width. **(b)** Representative images of the SLB (10% Brain PS + 89.5% Egg PC + 0.5% Texas-Red-PE) evenly wrapping around 300 nm width nanobars (magenta) with 5.4 μM bacterial expression eGFP-nsP1-(b) protein (green) on the bilayer. **(c)** Average images and the corresponding 3D surface plots of the lipid bilayer and eGFP-nsP1-(b) on nanobars of 300 nm width. n= 58 bars. **(d)** The end-to-center ratio of the SLB and eGFP-nsP1-(b) on nanobars. Data are the mean ±SD. P value calculated using unpaired t-test with Welch’s correction: **** represents p < 0.0001. n = 164 bars, with 3 independent experiments. **(e)** SEM image of the gradient nanobar structure with increasing width from 100 nm to 600 nm (two columns for each size, 100 nm step set). Individual bars with 200 nm, 400 nm, and 600 nm widths are shown in the lower panel. **(f)** Average images and corresponding 3D surface plots of the eGFP-nsP1-(b) on nanobars in width gradient from 100 nm to 600 nm over 80 bars. **(g)** Normalized nanobar-end density of eGFP-nsP1-(b) over 80 bars. Data are the mean ± SEM, with 3 independent experiments. **(h)** Representative images of confocal FRAP test for lipid bilayer (magenta) and eGFP-nsP1-(b) (green) on 200 nm width nanobar in 90 s. Yellow dashed circles indicate the bleaching area. **(i)** Normalized fluorescence intensity plot of lipid bilayer and eGFP- nsP1-(b) within nanobar area from FRAP measurement (2 s interval), from 3 independent experiments. **(j)** Representative images of the SLB (magenta), 1 μM Alexa Fluor 488 labeled nsP1 (green) on 99.5% POPC + 0.5% Texas-Red-PE SLB, 30% POPS + 69.5% POPC + 0.5% Texas-Red-PE SLB, 50% POPS + 49.5% POPC + 0.5% Texas-Red-PE SLB respectively on gradient nanobars with width from 100 to 600 nm. **(k)** Relative protein density of nsP1 for different lipid compositions over 80 bars. Data are the mean ±SD. P value calculated using unpaired t-test with Welch’s correction: **** represents p < 0.0001, with 3 independent experiments. **(l)** Normalized nanobar-end density of nsP1-(488)-(b) based on the corresponding lipid bilayer intensity on 30% POPS and 50%POPS SLB over 80 bars. Data are the mean ± SEM [standard error of the mean], with 3 independent experiments. Scale bars: 2 μm.

To examine the range of membrane curvature that the CHIKV nsP1 senses, we employed a nanobar array with the curvature gradient with the half-circle diameter ranging from 100 nm to 600 nm in 100 nm increments (SEM images shown in **Fig. 2e**). The purified eGFP-nsP1-(b) generated a stronger binding to smaller nanobars than the larger ones, as shown in both representative images (**Supplementary Fig. 6)** and the averaged ones (**Fig. 2f**). The protein density at differently sized nanobar ends was calculated by the ratio of nsP1 intensity at the nanobar end over the lipid fluorescence intensity per unit area. By plotting the normalized bar-end density over different nanobar diameters, the higher the membrane curvature generated, the higher the protein density obtained (**Fig. 2g**). Especially when the membrane was deformed with curvature greater than 300 nm diameter, the nsP1 demonstrated increasing curvature-dependent binding preference as the curvature diameter decreased. This suggested that during the formation of viral membrane spherule, the progressive increment of positive membrane curvature formed in the x-z plane of the saddle curvature enhanced the nsP1 accumulation on the membrane. However, different from the low FRAP recovery (∼20%) of mammalian-expressed nsP1 on live cell PM (**Fig. 1l, m**), the bacteria-expressed eGFP-nsP1-(b) showed much higher recovery (∼70%) on SLB-coated nanobars after photobleaching **(Fig. 2h, i, Supplementary Movie 3**), demonstrating higher mobility on the membrane. Considering the fact that no post-translational modification existed in the bacterial expression system, eGFP-nsP1-(b) used *in vitro* here lacked the S-palmitoylation at the cysteine residues. Earlier work reported that the S-palmitoylation directed nsP1 to cholesterol-rich regions on the cell membrane.^47^ Therefore, lacking both cholesterols in SLB and the S-palmitoylation on eGFP-nsP1-(b) here could lead to the observed mobility increase.

The impact of membrane charges on the curvature-dependent membrane binding of nsP1 was also evaluated. Recent studies showed that negatively charged lipids enhance the membrane binding of nsP1 via electrostatic interaction.^13,25,43^ However, whether or how it affects the curvature sensitivity of nsP1 is unknown. Here, we measured the nsP1 binding on nanobar-curved SLB containing negatively charged 16:0-18:1 phosphatidylserine (POPS) at different concentrations (0%, 30%, and 50%) (**Fig. 2j- l**). Consistent with prior studies using liposomes and giant unilamellar vesicles,^13,25^ nsP1 poorly bound to neutrally charged SLB with POPC only as evidenced by low fluorescent intensity across all the nanobar structures (**Fig. 2j**). With the increase of POPS from 30% to 50%, nsP1 showed higher fluorescence intensity both on nanobars and on the flat area in between (**Fig. 2j**), confirming the electrostatics-dependent membrane binding of the nsP1. The relative protein density measured on nanobars increased significantly from 0% POPS (0.079 ±0.036) to 30% POPS (0.448 ±0.145) and 50% POPS (0.642 ±0.192) (**Fig. 2k**). However, the normalized protein density at the nanobar end exhibited no significant difference across all the nanobar dimensions (**Fig. 2l**), indicating a minor influence of membrane charges across the range of curvatures that nsP1 could sense.

In addition to PS, we also examined another negatively charged lipid, 16:0-18:1 phosphatidic acid (PA) *in vitro* (**Supplementary Fig. 7a**), which was also anionic but with smaller head groups than PS. Similar to the effect of PS, higher PA concentration resulted in the increase of binding density of nsP1 on SLB (**Supplementary Fig. 7b**), but not the sensitivity to different curvature values (**Supplementary Fig. 7c**). However, comparing the results of PS and PA in the same concentration (30%), it was interesting to note that nsP1 enriched more on PA than PS containing SLB (**Supplementary Fig. 7d**). This was likely due to the fact that PA had smaller headgroup than PS and thus offered more lipid packing defects at the curved membrane for the curvature-sensitive protein to bind.^48^ Overall, our results here demonstrated that the geometry of lipid molecules had a stronger impact on the curvature sensitivity of nsP1 than their charges, although negatively charged lipids were necessary to ensure sufficient binding density of nsP1 on the membrane.

### 3. Hydrophobic residues of CHIKV nsP1 are essential for curvature sensing

To understand the curvature sensing mechanism of CHIKV nsP1, we further examined the role of several characteristic domains reported earlier for membrane binding in CHIKV or its closely related Semliki Forest virus (SFV) in the *alphavirus* genus. These included two recently identified membrane association (MA) loops around amino acid (a.a.) 220 to 233 (MA loop 1) and a.a. 407 to 427 (MA loop 2), a palmitoylation site at a.a. 417 to 419 near the C-terminal,^47^ and an amphipathic helix (AH) corresponding to a.a. 244 to 263^25^ that was previously demonstrated for effective membrane deformation *in vitro* and affecting the membrane association in cells for SFV **(Fig. 3a)**.^45,49^ It is to be noted that, although the AH of CHIKV nsP1 was recently revealed as non-critical for its membrane binding, it was still included here as an important control to exclude any potential influence of it on nsP1’s curvature sensing ability. This is because curvature sensing is not equal to membrane binding and can be altered via weaker molecular interactions^50^ than the stronger affinity needed for membrane binding. Accordingly, a series of mutations targeting these domains individually or combinatorically were constructed for evaluation, together with two truncations removing the flexible regions that have not been structurally resolved (a.a. 476-516, and a.a. 516-535)^45,49^ (illustrated in **Fig. 3b and Supplementary Fig. 12a**).

**Figure 3.**
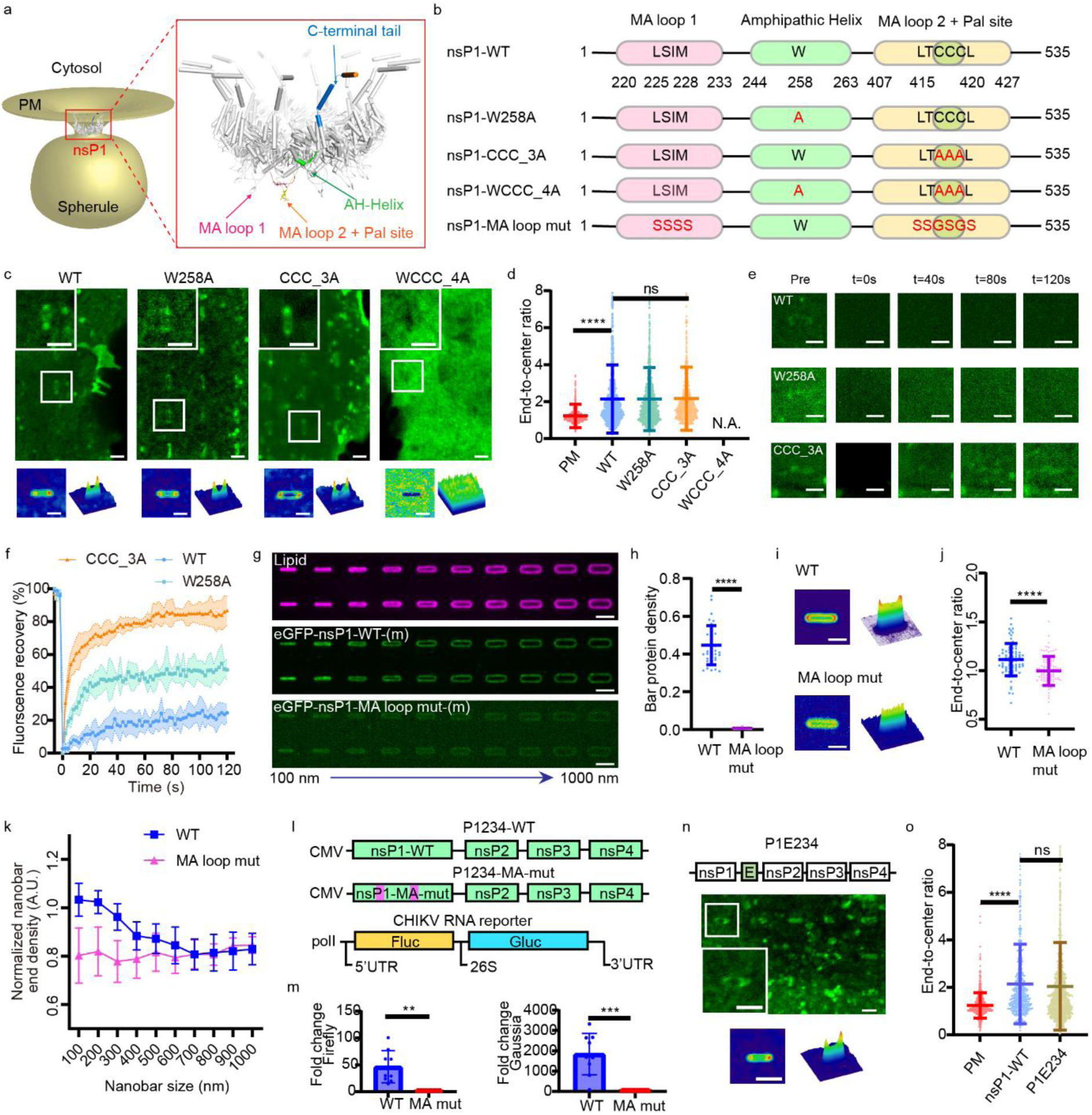
The mechanism of the CHIKV nsP1 curvature sensing ability. **(a)** A model of CHIKV nsP1 dodecamer identified the putative membrane binding domains on one of the twelve nsP1s: two MA loops (a.a. 220 to 233 and a.a. 407 to 427) with the palmitoylation site (a.a. 417 to 419) at MA loop 2, one AH (a.a. 244 to 263), and a disordered c-terminal tail (a.a. 474 to 535). **(b)** Mutations and truncations of CHIKV nsP1 used in this study. **(c)** Representative images of cells on nanobars expressing eGFP tagged nsP1-WT, nsP1-W258A, nsP1-CCC_3A, nsP1-WCCC_4A. The average images and corresponding 3D surface plots of the cells are shown below (n = 291, 763, 533, 424 bars respectively). **(d)** The end-to-center ratio of PM, nsP1-WT, nsP1-W258A, nsP1-CCC_3A, and nsP1- WCCC_4A on nanobars. Data are the mean ±SD. P value calculated using unpaired t-test with Welch’s correction: **** represents p < 0.0001. n =1027, 1250, 1255, 756 bars respectively, with 3 independent experiments. **(e)** Representative images of confocal FRAP test for nsP1-WT, nsP1-W258A, nsP1- CCC_3A in 120 s. **(f)** Normalized fluorescence intensity plot of nsP1-WT, nsP1-W258A, nsP1-CCC_3A within nanobar area from FRAP measurement (2 s interval). **(g)** Representative images of 30% POPS + 69.5% POPC + 0.5% Texas-Red-PE SLB (magenta) with 1 μM eGFP tagged mammalian expressed nsP1-WT (eGFP-nsP1-WT-(m)) or nsP1 MA loop mutant (eGFP-nsP1-MA loop mut-(m)) (green) on gradient nanobars with width from 100 to 1000 nm. **(h)** Relative protein density of nsP1- WT-(m) (n = 36 bars) and nsP1-MA loop mut-(m) (n = 75 bars). Data are the mean ± SD. P value calculated using unpaired t-test with Welch’s correction: **** represents p < 0.0001, with 3 independent experiments. **(i)** Average images and corresponding 3D surface plots of nsP1-WT-(m) and nsP1-MA loop mut-(m) on 300 nm width nanobars. n = 75 bars for WT and n = 93 bars for the mutant. **(j)** The end-to-center ratio of nsP1-WT-(m) and nsP1-MA loop mut-(m) on 300 nm width nanobars. Data are the mean ± SD. P value calculated using unpaired t-test with Welch’s correction: **** represents p < 0.0001. n = 75 bars for WT and n = 93 bars for the mutant, with 3 independent experiments. **(k)** Normalized nanobar-end density of nsP1-WT-(m) and nsP1-MA loop mut-(m) based on the corresponding lipid bilayer intensity over 70 bars. Data are the mean ±SEM, with 3 independent experiments. **(l)** Schematic representation of CMV-P1234, its MA loop mutant (CMV-P1234-MA-mut) and the RNA reporter (polI-Fluc-Gluc) constructs. CMV: immediate early promoter of human cytomegalovirus; polI: truncated promoter for human or Aedes albopictus RNA polymerase; Fluc: firefly luciferase; Gluc: *Gaussia* luciferase. **(m)** The RNA reporter expression level directed by genomic (firefly luciferase) and subgenomic (*Gaussia* luciferase) promoters after co-transfected with P1234 or P1234-MA-mut. Data are the mean ± SD. P value calculated using unpaired t-test with Welch’s correction: ** represents p < 0.01, *** represents p < 0.001. n = 9 for both conditions, with 3 independent experiments. **(n)** Representative images of cells on nanobars expressing eGFP tagged P1E234. The average images and corresponding 3D surface plots of the cells are shown below. n=1103 bars. **(o)** The end-to-center ratio of the PM, nsP1-WT, P1E234 on nanobars. Data are the mean ±SD. P value calculated using unpaired t-test with Welch’s correction: **** represents p < 0.0001. n = 666, 626, 1491 bars respectively, with 3 independent experiments. Scale bars: 2 μm.

Here, we firstly evaluated the influence of different domains in cells via transient transfection, which mainly focused on AH and palmitoylation, as the MA loops mutant was hardly expressed in U2OS cells and thus not possible to measure its association at nanobar-curved membrane sites (**Supplementary Fig. 8**). Specifically, AH domains have been recognized as typical curvature-sensitive domains widely used by many proteins, including epsin,^51^ α-Syn,^52,53^ Arf1,^54^ and ALPS motifs^53^. Even in alphavirus, SFV has been reported to contain an AH domain (a.a. 245 to 264) with a similar sequence as CHIKV (a.a. 244 to 263), and this AH has been shown to both interfere with the membrane-binding affinity of nsP1 in live cells and significantly altered membrane curvature *in vitro*.^20,47,55^ Although the same AH showed less impact on CHIKV membrane binding, whether it alters the sensitivity of nsP1 interacting with curved membranes indirectly is unknown. Here, we tested the AH mutation W258A (nsP1-W258A) which showed impaired membrane binding of nsP1 in SFV.^20^ When expressed in cells, nsP1-W258A generated preferential accumulation at the nanobar ends over the bar center (**Fig. 3c**), despite a decreased membrane binding than nsP1-WT (**Supplementary Fig. 9**) consistent with previous studies.^47^ The curvature enrichment was similar to the wildtype of nsP1 (nsP1-WT) with no significant difference in the measured end-to-center-ratio (WT: 2.143 ±1.840 *vs.* W258A: 2.143 ±1.710, unpaired t-test, p = 0.9953) (**Fig. 3d**). It indicated that the AH domain in nsP1 had indeed negligible influence on its curvature sensing ability. It was not surprising as the recent cryo-EM structure of CHIKV nsP1 revealed that the ‘AH’ (a.a. 244-263) of CHIKV nsP1 was an integral part of the protein core and less likely exposed to membrane for interactions (**Fig. 3a**),^45^ underscoring its minimal impact on curvature sensing abilities. These results also presented a non-conventional example that the predicted AH domains from protein sequence analysis might not always contribute to the curvature sensing depending on the actual protein conformation upon proper folding.

Besides AH, a palmitoylation-defective mutant at a.a. 417-419 (nsP1-CCC_3A) was also reported to decrease the membrane binding of nsP1 in CHIKV, and its combined double mutation with the AH W258A (nsP1-WCCC_4A) abolished the membrane fraction in cells.^47^ When tested on nanobar arrays here, the double mutant nsP1-WCCC_4A produced dominant cytosolic signals with no membrane fraction detectable on the nanobars (**Fig. 3c**), consistent with earlier reports.^47^ The CCC_3A mutant, however, displayed clear enrichment at nanobar ends (**Fig. 3c**) with a similar end-to-center ratio to the nsP1-WT (WT: 2.143 ±1.840 *vs.* CCC_3A: 2.159 ±1.702, unpaired t-test, p = 0.8367) (**Fig. 3d**), suggesting a non-essential role of palmitoylation for the curvature-guided distribution of nsP1 on the PM. In FRAP tests, nsP1-W258A exhibited no detectable recovery after photobleaching, while nsP1- CCC_3A recovered quickly (**Fig. 3e, f, Supplementary Movie 4**). It further supported that, although palmitoylation might be dispensable for the membrane anchorage of nsP1, it played a significant role in modulating its mobility on the PM. Overall, neither the AH nor the palmitoylation domain was the major determinant for the curvature sensing of nsP1 in CHIKV.

Besides AH and palmitoylation, the insertion of the protein’s hydrophobic residues has also been reported to contribute to membrane curvature sensing in various proteins, such as the C2 domains of Doc2b and synaptotagmin-1.^56,57^ For nsP1, the recent cryo-ET study identified two MA loops that contain 4 hydrophobic residues at a.a. 220 to 233 and 8 hydrophobic residues at a.a. 407 to 427 together with the palmitoylation site a.a. 417 to 419 (**Fig. 3a**).^45,49^ These MA domains are highly conserved among alphaviruses.^45^ Here, we evaluated the contribution of the hydrophobic residues from the MA loops on nsP1’s curvature sensing. For this purpose, we partially reduced the hydrophobic a.a. in MA loops 1 and 2, as shown in **Fig. 3b**, and obtained their purified protein from Expi293F cells (eGFP- nsP1-MA loop mut-(m) in short) for *in vitro* characterization. The negative-staining EM image showed that twelve copies of nsP1 MA loop mutant organized into C12 symmetry, same as the dodecamer ring observed for WT (**Supplementary Fig. 10a**).^45^ Gel-based MTase/GTase assay also confirmed the mRNA capping activity of the nsP1 MA loop mutant (**Supplementary Fig. 10b)**. The curvature sensing ability of the nsP1-MA loop mutant was then tested using the nanobar-based SLB assay *in vitro*. As shown in **Fig. 3g**, 1 μM eGFP-nsP1-WT-(m) or eGFP-nsP1-MA loop mut-(m) were incubated with 30% POPS SLB. The protein density of nsP1-MA loop mutant bound on nanobars was significantly decreased (0.006 ±0.003) compared with the nsP1-WT (0.447 ±0.103) (**Fig. 3h**), which confirmed that the MA loops were critical for anchoring nsP1 to membranes. Interestingly, there were two obvious peaks at the curved ends shown in the average image of mammalian expression nsP1 protein, but an even distribution of the mutant around the nanobar (**Fig. 3i**). The curvature sensitivity of the mutant (1.032 ±0.139) probed by the end-to-center ratio was significantly lower than the WT (1.134 ±0.186) at 300 nm wide nanobar (**Fig. 3j**). It suggested that the nsP1-MA loop mutant not only bound less to the membrane but was also less sensitive to curved membranes. Further probing their curvature response on gradient nanobar arrays showed that nsP1 WT exhibited increased protein density at nanobars with end curvature of 500 nm diameter or smaller, while nsP1-MA loop mutant showed much less preference for higher curvature at smaller nanobars (**Fig. 3k**). It proved that the hydrophobic residues in the MA loops were critical for nsP1 curvature sensing. Moreover, the impact of curvature-anchorage MA loops on the viral replication efficiency was further evaluated by employing a CHIKV *trans*-replication system which contained two plasmids to encode the P1234 polyprotein (or its MA loop mutant) and the RNA replication reporters separately (**Fig. 3l, Supplementary Fig. 11**). Here, the polyprotein P1234 could be cleaved into the 4 individual nsPs when expressed in cells. In addition, firefly and *Gaussia* luciferase genes were designed as reporters for genomic and subgenomic replication, respectively. Compared with the wild type nsP1 in P1234 (P1234-WT), the MA loop mutant (P1234-MA-mut) resulted in a significantly decreased expression level of replication reporters for both genomic and subgenomic region (**Fig. 3m**). It confirmed the critical role of the MA-loop in guaranteeing efficient viral replication.

In addition to hydrophobic insertion, intrinsically disordered regions have recently been shown as a new contributor to membrane curvature sensing.^58–62^ In nsP1, the entire C-terminal region (a.a. 474- 535) was reported to be disordered (**Fig. 3a**).^45,49^ To investigate whether it affected the curvature response, we truncated the a.a. from 477 to 535 at the C terminal tail (nsP1-(1-476), as shown in **Supplementary Fig. 12a**). Compared to the wildtype nsP1, nsP1-(1-476) exhibited a much stronger curvature sensing ability with higher end-to-center ratio (WT: 2.039 ±1.526 *vs.* 1-476: 3.011 ±3.235, unpaired t-test, p < 0.0001) (**Supplementary Fig. 12b**-d). However, a partial truncation of this disordered region, nsP1-(1-516) (**Supplementary Fig. 12a**) tuned the curvature response back to the wildtype level (WT: 2.039 ± 1.526 *vs.* 1-516: 2.055 ± 1.428, unpaired t-test, p = 0.8992) (**Supplementary Fig. 12b**-d). It indicated that residues 476 to 516 potentially hindered the nsP1 curvature sensing ability in live cells. One explanation of the decrease of the nsP1 curvature sensing ability when having a.a.477-516 in live cells was that negatively charged a.a were dominant between a.a. 477 to 516 (12 vs. 6 among 40 a.a., **Supplementary Fig. 13**) which was electrostatically repulsive towards the negatively charged lipid membrane. Another possibility was the steric hindrance of the disordered region which was less effective when detached from the nsP1.

Beyond the nsP1 protein itself, functional viral replication requires all four nsPs to assemble together to anchor the virus RNA for replication.^42^ Whether the binding of other nsPs affected the curvature-promoted nsP1 accumulation was also evaluated by expressing the CHIKV nsP polyprotein P1E234 (with eGFP fused at the c-terminal of nsP1, as illustrated in **Fig. 3n**) in cells. The result showed an accumulated green fluorescence signal on the nanobar ends (**Fig. 3n**), suggesting that the co- existence and assembly with nsP2, 3, and 4 in cells did not inhibit the curvature preference of nsP1. The measured end-to-center ratio of P1E234 (2.044 ±1.843) showed no significant difference compared to nsP1-eGFP alone (2.146 ± 1.675, unpaired t-test, p = 0.216) (**Fig. 3o**). Consistently, the cumulative frequency distribution of P1E234 was indistinguishable from the nsP1-WT curve (**Supplementary Fig. 14**). Hence, the results proved that nsP1 senses and accumulates on curved membrane sites regardless of the binding of other nsPs. On the other hand, other CHIKV nsPs were found to relocate to curved membranes via their association with nsP1. As shown in **Supplementary Fig. 15**, the sole expression of GFP-tagged nsP3 mainly resulted in fluorescence signals in the cytosol, while the co-expression with nsP1 (through P123E4, GFP fused to the c-terminal of nsP3) resulted in a detectable fraction of nsP3 at the PM.

### 4. Validation of curvature-facilitated stabilization of nsP1 via molecular dynamics simulations

To further elucidate the molecular basis of the membrane curvature sensing by nsP1, we performed a molecular dynamics simulation of nsP1 (a.a. 36 to 473) on a lipid bilayer containing 20% POPS and 80% POPC. All-atom and coarse-grained simulations were performed to study individual nsP1 molecules and their dodecamer, respectively (**Supplementary Fig. 16**). For a single copy of the nsP1 molecule on the planar bilayer (**Fig. 4a, Supplementary Fig. 17**), the MA loop 1 of nsP1 showed charge-dependent binding to the membrane mediated by its positively charged amino acid sidechains and the negatively charged head groups of POPS lipids. However, the MA loop 2 with more hydrophobic residues tended to move away from lipid headgroups, failing to form stable binding of individual nsP1 on the membrane. To eliminate the possibility that the initial position of the nsP1 might influence its membrane interaction, we also flipped the nsP1 at 90 degrees and 180 degrees. The result consistently revealed that nsP1 did not form stable binding with the membrane in 20 ns under either of these configurations (**Supplementary Fig. 18**). In contrast, nsP1 dodecamer showed significantly stabilized binding to the membrane when introduced to the coarse-grained model (**Fig. 4b)**. After back- mapping^63^ the entire nsP1 dodecamer to its all-atom form, we clearly confirmed the bind was through the sequential insertion of nsP1 MA loop 2 into the planar membrane (**Fig. 4b, zoom-in area in the black box**). Furthermore, the binding of nsP1 dodecamer generated local positive membrane curvature on the initially flat membrane surface (**Supplementary Movie 5**). As clearly demonstrated in the heat- map plot of the mean membrane curvature (**Fig. 4c**), positive membrane curvature was generated at the site of MA loop binding and insertion (red-colored region). Such a variation in the membrane curvature was also clearly seen from the cutaway view in the final image of **Fig. 4b**.

**Figure 4.**
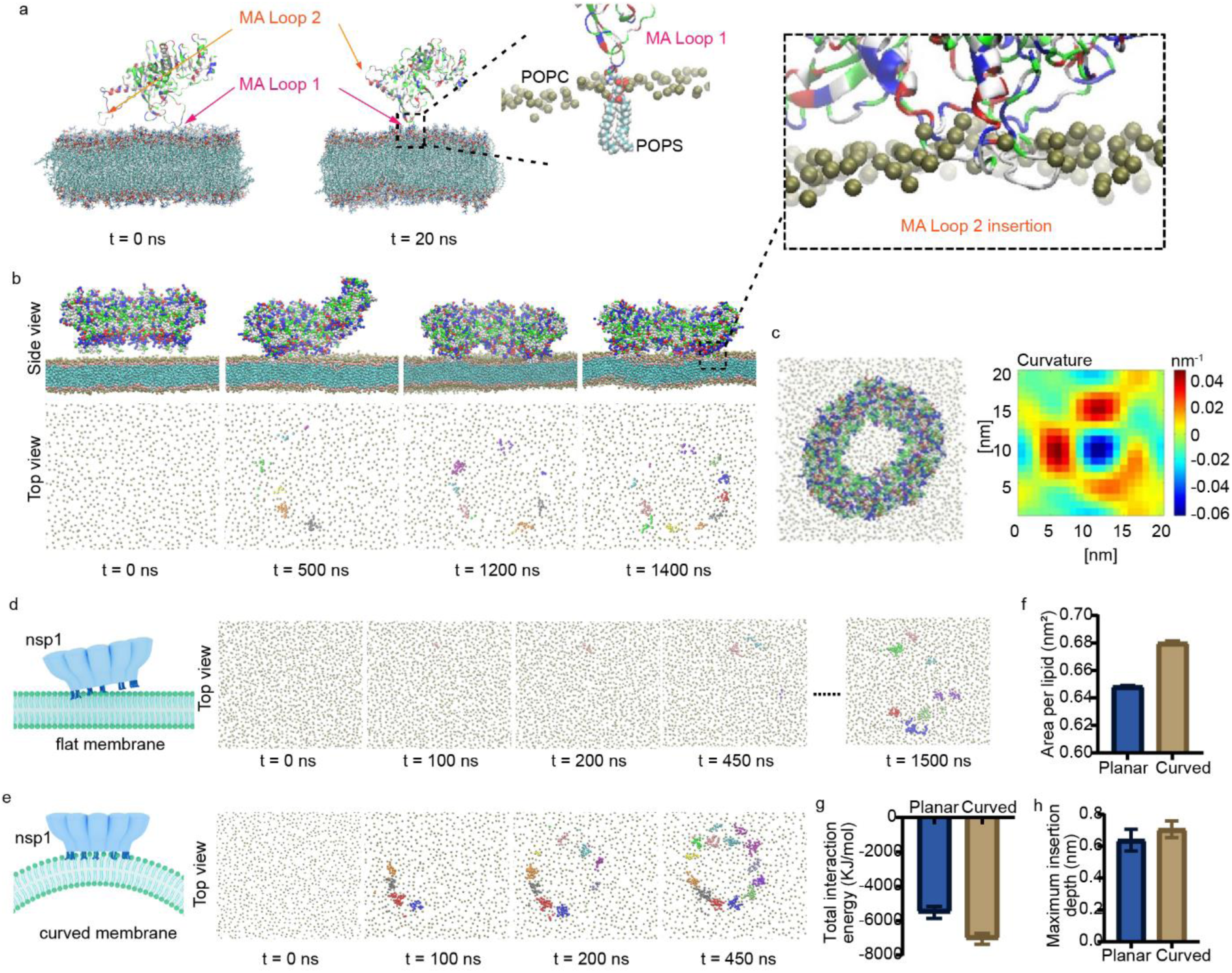
Molecular dynamic (MD) simulations of nsP1 protein-lipid membrane interaction. **(a)** Snapshots of all-atom MD simulation showing the interaction of a single copy of nsP1 protein with a planar membrane at initial (left) and equilibrated (right) states. Protein atoms are rendered based on the nature of residues (acidic: red; basic: blue; hydrophobic: white; and polar: green) while the atoms in lipids including carbon (C), oxygen (O), phosphorus (P), nitrogen (N) and hydrogen (H) are rendered in cyan, red, green, blue, and white, respectively. **(b)** Simulation snapshots showing the temporal evolution of the interaction of the nsP1 dodecamer with a planar membrane. The top panels show the side cutaway views of nsP1 dodecamer-membrane interaction at different simulation times while the bottom panels highlight the number of nsP1 ring sub-units inserted into the membrane at the same time. The lipid head beads (PO4, NC3, and CNO) are colored in tan, glycerol linkages (GL1 and GL2) in pink, and lipid tail beads in cyan, and only PO4 beads are shown in the top views. Protein beads in the top panels are rendered in the same way as in panel a while the inserted protein beads shown in the bottom panels are rendered in such a way that each color refers to a particular sub-unit of the dodecamer structure. Zoomed-in view of the region within the rectangular box highlights the insertion of nsP1 MA loop 2 into the planar membrane. **(c)** Top view of the planar membrane with nsP1 dodecamer bound (left) and the corresponding color map for mean membrane curvature (right). **(d)-(e)** Simulation snapshots showing the temporal evolution of the interaction of the nsP1 dodecamer with a flat supported membrane **(d)** and supported membrane with a radius of curvature of 100 nm **(e)**. **(f)** Comparison in the area per lipid between planar and curved membranes. **(g)** Comparison in the total interaction energy of nsP1 dodecamer with planar and curved membranes. **(h)** Comparison in the maximum insertion depth of nsP1 subunits (averaged over all inserted sub-units) into planar and curved membranes.

The generation of a positive curvature in the unrestricted membrane at the binding sites of nsP1 dodecamer suggested a more stable binding of nsP1 dodecamer to a positively curved membrane. To further test this speculation, we simulated the interaction of nsP1 dodecamer with SLBs with and without curvature (**Fig. 4d, e**). Although the nsP1 dodecamer could bind to both SLBs through the sequential insertion of hydrophobic loops into the lipid membrane, its binding to the curved membrane (**Fig. 4e**) was much more efficient than the flat one (**Fig. 4d**). While complete binding of the nsP1 ring to the curved membrane occurred within 450 ns of a production run, the complete binding to the flat membrane was not achieved even after 1500 ns. The accelerated binding of nsP1 dodecamer to the curved membrane was possibly due to the curvature-induced increase in area per lipid (**Fig. 4f**),^64^ which increased the appearance of the lipid packing defects and consequently the exposure of hydrophobic lipid tails to hydrophobic MA loops of nsP1. As expected, the nsP1 dodecamer bound stronger to the curved membrane as indicated by the more negative total interaction energy due to electrostatic and van der Waals interactions between nsP1 dodecamer and lipid membrane (**Fig. 4g**). The stronger binding between nsP1 dodecamer and the curved membrane was not only due to the higher number of MA loops inserted but also because the MA loops inserted deeper into the curved membrane (**Fig. 4h**). These simulation results proved that positive membrane curvature enhanced the nsP1 assembly on the membrane by facilitating the insertion of hydrophobic MA loop 2 and speeding up the binding kinetics, which was in good agreement with nanobar curvature-enhanced nsP1 stabilization on the cell PM.

### 5. Nanostructure enriched production of CHIKV viral genome

An exciting extension of the curvature-facilitated assembly of CHIKV nsPs for replication is the possibility of guiding the viral RNA genome replication on nanostructure arrays. Recent studies indicated that the formation of membrane spherule during successful viral RNA replication was to compensate for the pressure generated by the growing RNA polymers within the confined membrane compartments ^13^, which suggested that sufficient space was required near the replication sites to accommodate the membrane spherule growth without high pressure accumulated. In this sense, the nanobar design might not provide enough space nearby to allow the relaxation of the membrane tension built up during replication. To address this issue, we designed nanoring arrays where positive curvature could be generated in three places (as shown in **Fig. 5a-b**): (i) the sidewall of the outer ring, similar to the positive curve generated on nanobar with almost no space underneath the positively curved membrane; (ii) the top rim of the outer ring with sharper curvature than the side wall and similarly no space under it; and (iii) the top rim of the inner ring with sharper curvature similarly to the outer rim but extensive space close by and allowing more extension towards the center of the inner rings. When culturing cells infected with CHIKV on the nanoring arrays, we observed significant enrichment of double-stranded RNA (dsRNA) replication (**Fig. 5c, Supplementary Fig. 19**), together with nsP1 and nsP3 (**Fig. 5d**), in the center of the nanoring. In contrast, the replication sites of infected cells on flat surfaces were more randomly distributed throughout the PM (**Fig. 5c, Supplementary Fig. 19**). By measuring the dsRNA signal per unit membrane area marked by PM staining, we found that the dsRNA density inside each ring showed a significant elevation in contrast to the nearby flat area (**Fig. 5e)**. By averaging nanorings with a gradient of inner ring diameters (from 1500 nm to 700 nm in diameter with over 30 rings each), we observed a significant accumulation of dsRNA within the inner ring space, with higher dsRNA enrichment in larger rings. (**Fig. 5f**) Two possible factors might contribute to this correlation with inner ring diameter: the longer rim of the positively curved inner ring that could be measured by the inner rim perimeter; and the larger inner ring space proportional to the inner ring area. The contribution of the curved membrane and the nearby space could be compared by normalizing the dsRNA intensity with the inner ring area and perimeter respectively (**Fig. 5g, h**). Interestingly, a positive correlation between dsRNA accumulation and inner ring diameter was observed in both cases, which indicated that both the existence of the curved membrane and the availability of nearby space are beneficial for viral RNA replication.

**Figure 5.**
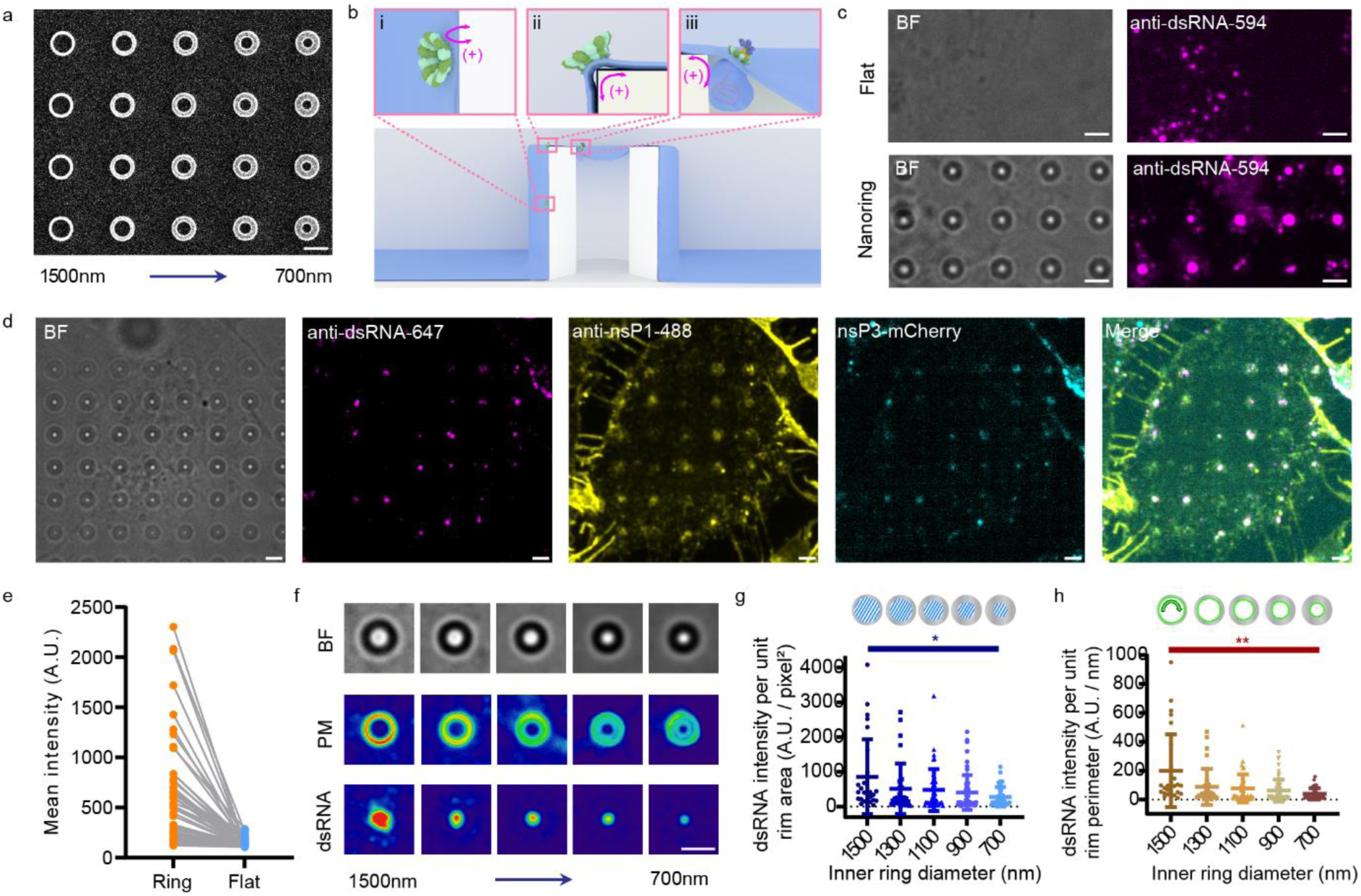
Nanostructure guided CHIKV replication. **(a)** An SEM image of gradient nanorings with decreasing inner ring diameter, ranging from 1500 nm to 700 nm. **(b)** A schematic representation of a nanoring illustrated three distinct positive curvatures on PM: (i) the sidewall of the outer ring; (ii) the top rim of the outer ring; and (iii) the top rim of the inner ring. **(c)** Representative images of CHIKV-infected cells cultured on a flat surface or a nanoring array. Replication sites are indicated by dsRNA staining (magenta). **(d)** Representative images of nsP3-mCherry tagged CHIKV-infected cells cultured on the nanoring array. The mCherry fluorescent tag was inserted at the hypervariable region of nsP3 (as shown in cyan) which is known to tolerate tags and does not impede virus replication and infection. The cells were also stained for nsP1 (yellow) and dsRNA (magenta). **(e)** Comparison of the dsRNA density in the inner ring of the nanoring area and on the nearby flat surface. n = 60 rings for both conditions, with 3 independent experiments. **(f)** Average images of the brightfield, PM intensity, and dsRNA intensity on different-sized nanorings (over 30 rings for each size). **(g)** The dsRNA intensity per unit inner rim area on different-sized nanorings. Data are the mean ±SD. P value calculated using unpaired t-test with Welch’s correction: * represents p < 0.05. n = 27, 31, 44, 60, 39 rings respectively, with 3 independent experiments. **(h)** The dsRNA intensity per unit inner rim perimeter on different-sized nanorings. Data are the mean ±SD. P value calculated using unpaired t-test with Welch’s correction: ** represents p < 0.01. n = 27, 31, 44, 60, 39 rings respectively, with 3 independent experiments. Scale bars: 2 μm.

The detailed arrangement of viral RC spherules on nanorings was further visualized in high resolution using expansion microscopy (**Fig. 6a, Supplementary Fig. 20, Supplementary Movie 6**). As shown in **Fig. 6b, c**, although the curvature-enhanced distribution of nsP1 appeared both surrounding the nanoring outer sidewalls and following the rims as expected, the successful viral genome replication marked by dsRNA staining exhibited dominant accumulation only next to the inner ring rim. A zoom-in view of the inner ring rim (**Fig. 6c, Supplementary Fig. 21**) further revealed that the dsRNA signals protruded towards the center space of the inner ring while the nsP1 located at the edge of the dsRNA puncta next to the nanoring rim marked by gelatin staining. It confirmed that both curvature-guided nsP1 distribution and nearby space were important for productive viral replication. In addition to the center space of the rings, the bottom of the outer ring also created available space for membrane spherule generation when cells were not wrapping the nanoring too tightly, and thus also allowed productive RNA replication as marked by dsRNA staining (**Fig. 6e**). Further zooming in on individual dsRNA puncta confirmed that there were nsP1s located at their edges and attached to the positively curved outer ring side wall as well (**Fig. 6f, Supplementary Fig. 22**). Interestingly, the bottom enrichment of dsRNA was only observed in samples with poor gelatin coating around nanorings (**Fig. 6d**) but was absent in samples with tighter gelatin coating where no space was detected at the bottom of the nanorings. It further validated that sufficient space was needed to allow membrane spherule generation for viral RNA replication.

**Figure 6.**
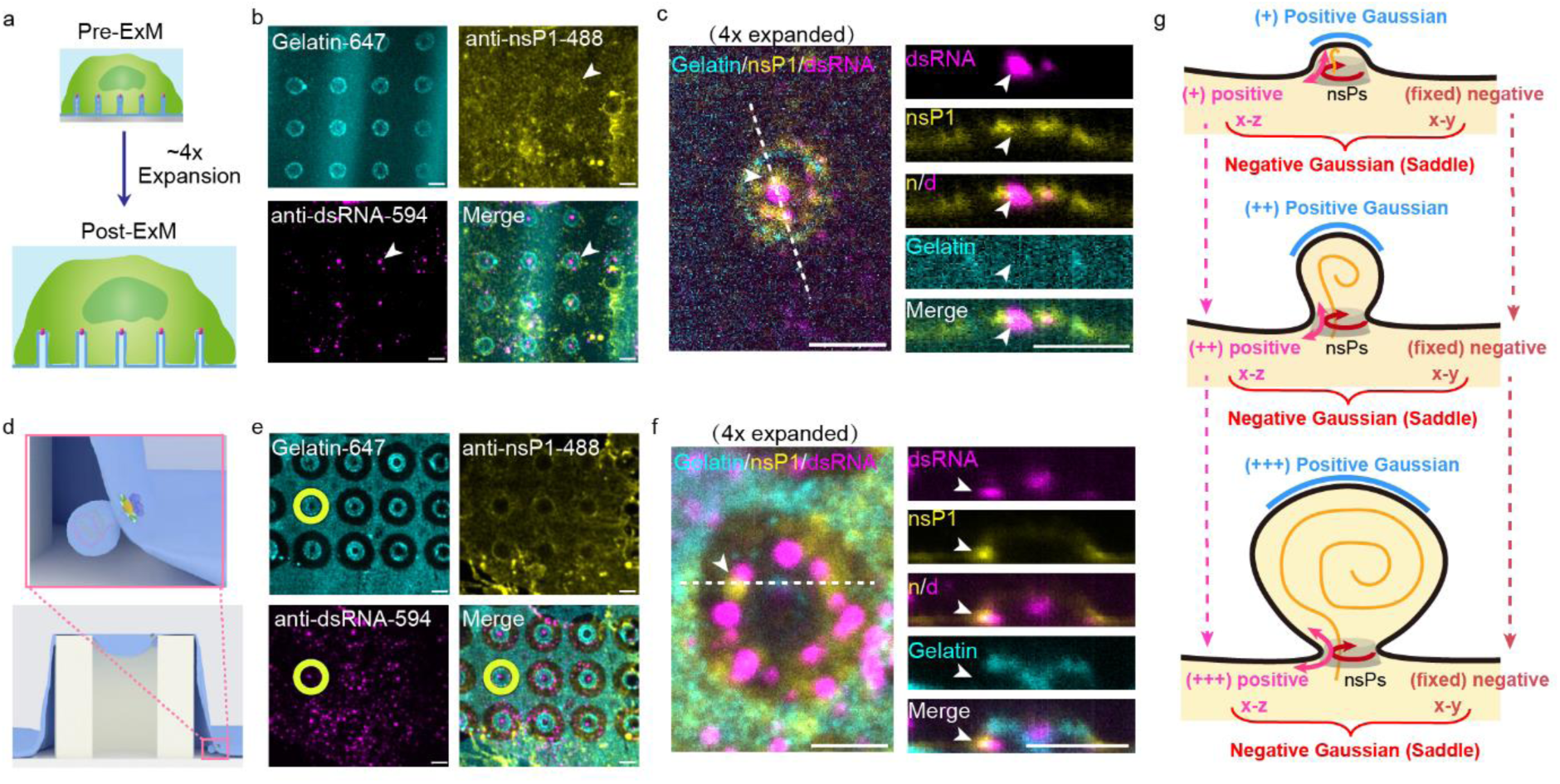
The detailed arrangement of viral replication complex on nanorings revealed by super- resolution imaging. **(a)** A schematic illustration of expansion microscopy (ExM) for higher-resolution imaging. **(b)** Post-ExM images of a CHIKV-infected cell cultured on fluorescent gelatin-coated nanorings (cyan). The cell was stained for nsP1 (yellow) and dsRNA (magenta). A representative replication site is indicated by the white arrow. **(c)** Zoomed-in x-y image (left) and the resliced z-axis image along the white dash line (right) both show one nanoring with uniform gelatin coating (cyan). The CHIKV-infected cell cultured on the nanoring surface exhibits dsRNA signals concentrated in the center region of the inner ring (magenta) while the nsP1 is located at the periphery of the dsRNA puncta next to the nanoring rim (yellow). **(d)** A schematic of nanoring shows that the incomplete PM adhesion to the nanoring sidewall can create a space at the bottom of the outer ring, allowing for CHIKV replication in this region. **(e)** Post-ExM images of a CHIKV-infected cell cultured on fluorescent gelatin-coated nanorings (cyan). Incomplete PM adhesion, indicated by dark voids of the gelatin coating around the nanoring (one example is highlighted as a yellow ring), allowed for viral replication within these spaces. The cell was stained for nsP1 (yellow) and dsRNA (magenta). **(f)** Zoomed-in x-y image (left) and the resliced z-axis image along the white dash line (right) both show the replication taking place at the bottom of the outer ring. **(g)** An illustration shows two distinct curvatures generated during spherule formation: 1) a saddle curvature at the neck of the spherule with an increasing positive curvature along the x-z plane (pink color) sensed by nsP1 due to the hydrophobic insertion of the MA loops, and a fixed negative curvature along the x-y plane (red color) fixed by the size of the nsP1 dodecamer ring; and 2) a progressively increasing positive curvature along the crown of the spherule (blue color). Scale bars: 2 μm.

## Conclusion

In summary, by designing nanostructures to decouple the saddle curvature into separated positive and negative curvatures, we revealed that nsP1 formulated an efficient viral protein system to modulate saddle curvatures formed on CHIKV membrane spherules for viral genome replication. As proved both in live cells and *in vitro*, individual nsP1 preferentially interacted with positively but not negatively curved membranes via the hydrophobic residues of the two MA loops. However, by forming a rigid ring, the nsP1 dodecamer not only bound to the membrane much more stable than the monomers but also generated a negative curvature naturally with a fixed diameter in the x-y plane that was perpendicular to the x-z plane with positive curvature for MA-loop insertions. Interestingly, as illustrated in **Fig. 6g**, the MA loops of individual nsP1 could sense a wide range of positive curvatures generated at the x-z plane of the membrane spherule neck, which indeed underwent a progressive increase during viral replication.^13,14^ On the other hand, the dodecamer ring of nsP1 posed a fixed negative curvature at the x-y plane of the neck to prevent fluctuation of the channel opening and relate transport regulation during the enlargement of the spherules along with viral genome replication, which also offered a stable anchorage of nsP2, nsP3, and nsP4 to the center of the ring-opening regardless the replication-induced positive curvature alterations in the x-z plane.^13,65,66^ Distinct from the saddle curvature sensing proteins and peptides reported earlier,^35,67–71^ the saddle curvature modulation of nsP1 decoupled the monomer-based sensing of positive curvature from oligomer-based generation of negative curvatures into two perpendicular planes. Moreover, our study also demonstrated a potential functional correlation between viral replication and membrane curvature adaptation, where curved membrane sites on PM could effectively guide the spatial distribution of CHIKV replication thanks to the curvature sensitivity of nsP1.

## Acknowledgments

We thank Dr. Andres Merits (University of Tartu, Estonia) for providing anti-nsP1 antibody, CHIKV P1234 plasmid, and HSPolI-Fluc-Gluc plasmid; Dr. Yee-Song Law (Lee Kong Chian School of Medicine, NTU) for constructing eGFP-nsP1 plasmid and purifying bacterial expressed nsP1 proteins; Ms. Sin Yi Chong (School of Chemistry, Chemical Engineering and Biotechnology, NTU) for maintaining cells during the early stage of this project. We thank the Nanyang NanoFabrication Centre (N2FC) and the Centre for Disruptive Photonic Technologies (CDPT) at Nanyang Technological University (NTU) for supporting nanostructure fabrication and SEM imaging; the School of Chemistry, Chemical Engineering and Biotechnology NTU and Dr. Yansong Miao (School of Biological Science, NTU) for generous support on the laser scanning and the spinning disk confocal microscopes. We thank the financial support provided by the China Scholarship Council (CSC) and the guidance by Dr. Junxian Yun (College of Chemical Engineering, Zhejiang University of Technology) for Dr. Jiawei Wu’s work. This work was supported by funding from the Singapore Ministry of Education under its MOE Academic Research Fund Tier 3 Award MOET32020-0001 and MOET32023-0004 (to W.Z.), Tier 2 Award MOE-T2EP30222-0022 (to W.Z.), MOE-T2EP30220-0009 (to D.L.), MOE-T2EP30124-0023 (to D.L.), MOE-T2EP50121-0004 (to C.H.) and Tier 1 grant RG112/20 (to W.Z.), RG95/21 (to W.Z.), RG93/22 (to W.Z.), RG84/21 (to D.L.), RT22/23 (to D.L.); National Research Foundation under NRF-MSG-2023-0001 (to W.Z.) and NRF/ASTAR quantum engineering programme NRF2021-QEP2-03- P10 (to W.G.); Human Frontier Science Program under Early-Career Research Grant (RGY0088/2021) (to W.Z.). We would also like to acknowledge the support received from the Ministry of Education, Singapore, under its Research Centre of Excellence award to the Institute for Digital Molecular Analytics (IDMxS, grant: EDUN C-33-18-279-V12) (to W.Z.), and Nanyang Technological University for its Start-Up Grant (to W.Z.). The computational work for this article was fully performed on resources of the National Supercomputing Centre, Singapore (https://www.nscc.sg).

## Author contributions Statement

X.M., D.L., and W.Z. conceived the study and designed the experiment. X.M. performed most of the experiments. M.L. purified the mammalian expressed nsP1 proteins and conducted the virus infection experiment. J.K., C.C., and C.H. performed the molecular simulation analysis. Y.P.Z., X.G., Y.Y.Z., and W.G. fabricated the nanostructures, performed SEM imaging, and tested the chips. M.L. and Y.T. performed TEM measurements. X.M. and J.W. analyzed most of the data. X.M. and L.H. performed the expansion microscopy. X.M. and W.Z. drafted the manuscript. All the authors discussed the results and commented on the manuscript. Project administration and funding acquisition: W.Z., D.L., C.H., and W.G.

## Method

### 1. Nanostructure chip fabrication

Nanostructures used in the project were fabricated on the quartz wafer (1.5*1.5 cm) by electron-beam lithography (EBL) technique. The quartz chips were firstly spin-coated with 300 nm positive electron-beam resist PMMA (Allresist), followed by one layer of conductive polymer, AR-PC 5090.02 (Allresist). Customized patterns were exposed by EBL (FEI Helios NanoLab). After that, isopropanol (IPA): methylisobutylketone solution (MIBK) = 3:1 solution was used to remove the exposed PMMA. Subsequently, a 30 nm chromium (Cr) mask was generated via thermal evaporation (UNIVEX 250 Benchtop) and lift-off in acetone. Nanostructures were finally synthesized through reactive ion etching (RIE) with a mixture of CF4 and CHF3 (Oxford Plasmalab 80). SEM (FEI Helios NanoLab) imaging was performed after 10 nm chromium coating to measure the dimensional properties of nanostructure arrays. Before use, the nanochips were cleaned with Chromium Etchant (Sigma-Aldrich) to remove the remaining metal layer.

### 2. Plasmids

CHIKV nsP1 gene was cloned into the vector pEGFP-C1 (Clonetech) following the eGFP gene with a tag composed of strep and the 3XFLAG tag was inserted after a.a. 516 residue, referred to as eGFP-nsP1-WT. CHIKV nsP3, P1234-WT, P1E234-WT, P123E4-WT, and RNA reporter (HSPolI-Fluc-Gluc) plasmids were received as generous gifts from Andres Merits.^72^ VEEV nsP1 gene was cloned into the same vector as CHIKV nsP1. For the generation of the mutants including CHIKV nsP1- W258A, nsP1-CCC_3A, nsP1-WCCC_4A, nsP1-MA loop mut, P1234-MA-mut, primers were designed to target amino acid residues for mutagenesis. In the case of the plasmids encoding truncated nsP1-(1-476) and (1-516), two stop codons were introduced after residue 476 and residue 516 respectively. Site-directed mutagenesis was done using the KOD FX Neo polymerase chain reaction kit (TOYOBO) on a thermal cycler. The PCR products were digested with DpnI restriction enzyme and transformed into DH5-alpha competent E. Coli cells (New England Biolabs). Transformed cells were plated on kanamycin 50µg/ml containing agar plates and incubated overnight at 37°C. Colonies were selected to grow in LB containing kanamycin 50 µg/ml. Plasmids were recovered with miniprep kit (Axygen) and successful mutants were identified via Sanger sequencing which were subsequently used in downstream experiments.

### 3. Cell culture on nanostructure array

*Homo sapiens* bone osteosarcoma U2OS cells (ATCC) were maintained in DMEM with GlutaMAX (Gibco) supplemented medium with 10% fetal bovine serum (FBS) (Life Technologies) and 1% Penicillin-Streptomycin (PS) (Life Technologies). One day before transfection or virus infection, cells needed to be cultured onto the nanochip. Before introducing cells to the chip, the surface of the chip needed to be cleaned thoroughly and then underwent high-power Air-plasma (Harrick Plasma) treatment for 3 minutes. After that, the chip was coated with 0.2% gelatin (Sigma-Aldrich) for 30 minutes to facilitate optimal cell adhesion. After coating, the cells were plated onto a 35mm dish with the nanochip attached to the bottom and waited overnight before transfection or virus infection. The cells were maintained under 37 ℃ with 5% CO2.

### 4. Cell transfection and membrane staining

Plasmids were transfected with 1 μg plasmid mixed with 1.5 μL lipofectamine 3000 (Life Technologies), 2 μL P3000 reagent (Life Technologies), and Opti-MEM (Gibco) and incubated for 30 minutes. The cells were then starved in Opti-MEM with the plasmid mixture for 4 hours, after which they were incubated for 18 hours in the DMEM-supplemented medium with 10% FBS and 1% PS to promote protein expression for imaging. CellMask Deep Red (Life Technologies) staining was performed as a membrane control before imaging. The 1000-time diluted dye was added to the cells and incubated for 2 minutes at 37 ℃ with 5% CO2. The dye was then washed with the culture medium to prevent its intake inside the cells.

### 5. Protein purification

The method for preparing bacterial cell expressed CHIKV nsP1 was described in Zhang et al.^45^ Monomer fraction (peak 3) was collected and subsequently labeled with Alexa 488 using Alexa Fluor™ 488 Protein Labeling Kit (Invitrogen™) as per manufacturer’s instructions. For the mammalian expression system, plasmids were transfected into Expi293F cells by PEI MAX® (Polysciences) for transient expression, protein expression was boosted by 10 mM sodium butyrate 16 to 20 hours after transfection. Cell pellets were harvested after 96 hours and lysed in lysis buffer 50mM Tris-HCl pH 8.0 150mM NaCl 0.5mM TCEP with protease inhibitor, 1% n-dodecyl-β-D-maltoside (DDM), and 70 mU/ml of BioLock solution (IBA Lifesciences). The pellet was incubated at 4°C for 2 hours on a rotator with an occasional vortex to ensure complete suspension of the pellet in the lysis buffer. It was subsequently sonicated for 10 minutes, and centrifuged for 1h at 18,000g at 4°C. The supernatant was applied to an Econo-Pac® chromatography column (Biorad) containing Strep-Tactin® Sepharose beads (IBA Lifesciences). After serial washing with 5, 25, and 50 column volumes with the wash buffer (50 mM Tris-HCl pH 8.0, 150 mM NaCl, 0.5 mM TCEP, 10% glycerol), the proteins were eluted using wash buffer with added 10 mM D-biotin (IBA Lifesciences). The elutants were further purified through Superose® 6 increased 10/300 GL (GE Healthcare Life Sciences) or concentrated to the desired concentration for downstream experiments.

### 6. Liposome preparation

Lipid to test nsP1 curvature sensing consisted of 89.5 mol% Egg PC (L-α-phosphatidylcholine (Egg, Chicken), Avanti), 0.5 mol% Texas-Red-PE (Texas Red™ 1,2-Dihexadecanoyl-sn-Glycero-3- Phosphoethanolamine, Triethylammonium Salt, Life Technologies) and 10 mol% Brain PS (L-α- phosphatidylserine (Brain, Porcine) (sodium salt), Avanti). Lipid to study the POPS lipid charge effect on nsP1 curvature sensing consisted of POPC (1-palmitoyl-2-oleoyl-glycero-3-phosphocholine, Avanti), 0.5 mol% Texas-Red-PE with increasing mol% POPS (1-palmitoyl-2-oleoyl-sn-glycero-3- phospho-L-serine (sodium salt)) from 0 mol%, 30 mol% to 50 mol%. Lipid to study the PA lipid charge effect on nsP1 curvature sensing consisted of POPC, 0.5 mol% Texas-Red-PE with increasing mol% PA (1-palmitoyl-2-oleoyl-sn-glycero-3-phosphate (sodium salt)) from 0 mol%, 10 mol% to 30 mol%. To study the MA-Loops effect on eGFP-nsP1-(m) curvature sensing, the lipid consisting of 69.5 mol% POPC, 0.5 mol% Texas-Red-PE, and 30 mol% POPS was used.

For liposome formation, the lipid mixture with the desired composition was dissolved in chloroform and dried in a brown glass vial under 99.9% nitrogen gas for 5 minutes, followed by vacuum drying for at least 3 hours to remove the remaining chloroform. The dried mixture was then resuspended in PBS (Phosphate Buffered Saline, Gibco) buffer to achieve a concentration of 2 μg/μL and sonicated for 30 minutes. This was followed by a freeze-thaw cycle performed 20 times (20 seconds in liquid nitrogen, 2 minutes in a 42°C water bath) until the lipid mixture became clear. Subsequently, the mixture was extruded using an extruder equipped with a holder/heating block (610000-1EA, Sigma-Aldrich), passing through a 100 nm pore-size polycarbonate membrane 20 times until the solution appeared clearer. The resulting lipid vesicle solution was collected and stored at 4°C in preparation for SLB formation.

### 7. Supported lipid bilayer formation on nanochips

The nanochips were firstly cleaned with piranha solution (1 part 30% hydrogen peroxide solution and 7 parts concentrated sulfuric acid) overnight. Afterward, the chips were rinsed with a continuous stream of deionized water, dried with 0.45 μm-filtered 99.9% nitrogen gas, and hydrophilized and cleaned with high-power Air-plasma (Harrick Plasma) treatment for 1 hour. Lipid vesicles were then loaded onto the well-cleaned nanochips. The chips were attached to a polydimethylsiloxane (PDMS) chamber to prevent the lipids from drying. After a 15-minute incubation, excess lipid vesicles were washed away with 200 μL PBS buffer five times to form the lipid bilayer. Subsequently, the protein solution was added to the lipid-bilayer-coated nanochips, incubated for 15 minutes, and prepared for fluorescence imaging.

### 8. Cryo-EM sample preparation and microscopy

Briefly, copper grids (Carbon Film 300 Mesh Cu, Electron Microscopy Sciences) were glow discharged for 45 seconds and prepared with 10 μl of purified protein sample. After 1 min incubation, the excess sample was removed with filter paper and 2% uranyl acetate stain (Electron Microscopy Sciences) was applied for 1 min with excess stain removed with filter paper. Negative staining samples were screened on ThermoFisher FEI Tecnai T12 microscope at 120 kV with Eagle camera.

### 9. RNA 5’ end capping assay

The method previously described by Li et al.^73^ and optimized for CHIKV nsP1 in Law et al.^74^ CHIKV 5′ UTR sequence was used as the template to synthesize the substrate RNA. The 5′ triphosphorylated (ppp) single-stranded 12-mer RNA AUGGCUGCGUGA labeled with a FAM dye was named “pppAU-10 FAM”. A 5′ diphosphorylated (pp) single-stranded 12-mer RNA AUGGCUGCGUGA was labeled with a FAM dye at the 3′ end and named as “ppAU-10 FAM”. A Cap-0 RNA was synthesized based on the above sequence, named “m^7^GpppAU-10” or labeled with FAM dye was named “m7GpppAU-10 FAM” respectively. In addition, 5′ triphosphorylated (ppp) single-stranded 12-mer RNA AGUUGUUAGUCU was based on the dengue virus 5′ UTR labeled with FAM dye was named “pppAG-10 FAM” and a Cap-0 RNA “m^7^GpppAG-10 FAM”. The RNAs were synthesized by Trilink Biotechnologies and Bio-Synthesis Inc. The capping reaction was in two steps. Firstly, a covalent nsP1-m7GMP intermediate reaction was prepared in a 20 μL containing 20 μM nsP1, 50 mM Tris-HCl pH 7.5, 2 mM DTT, 10 mM KCl, and 2 mM MgCl2 and 0.5 mM S-adenosylmethionine (SAM) and 1 mM GTP or an alternative substrate as cap donor was incubated at 30°C for 2 h. Secondly, after the preparation of the covalent m7GMP-nsP1 was used for the transfer to RNA recipient in a 20 μL mixture containing 5 μL covalent intermediate, 50 mM Tris-HCl pH 7.5, 2 mM DTT, 2 mM MgCl2, and 1 μM synthetic RNA, and 20 U Murine RNase Inhibitor (New England Biolabs). The reactions were incubated at 30°C for 12 h and terminated by adding the stop solution 2X RNA loading dye (95% formamide 0.02% bromophenol blue 0.01% xylene cyanol 0.02% SDS 1 mM EDTA). The capped RNA products were separated by 20% denaturing RNA 8M urea PAGE gel and visualized using ChemiDoc™ MP imaging system (Biorad).

### 10. The *trans*-replicase system for analyzing CHIKV replicase activities

HEK293FT cells were maintained in DMEM with GlutaMAX supplemented medium with 10% FBS and 1% PS under 37 ℃ with 5% CO2. Cells grown on 24-well plates were co-transfected with 0.5 μg of plasmid encoding RNA reporter (HSPolI-Fluc-Gluc) with 0.5 μg P1234 or P1234-MA-mut by PEI MAX® (Polysciences). After 24 hours of incubation, the cells were washed with PBS and lysed. The expression levels of firefly and Gaussia luciferase were measured using the Dual-Luciferase® Reporter Assay System (Promega) on a 96-well plate (Corning) with a microplate reader (Synergy™ H1, BioTek). The activities of the RNA reporters were normalized to control cells transfected with the sole RNA reporter.

### 11. CHIKV infection in cells

U2OS cells were cultured on the nanochip one day before the CHIKV infection. On the day of infection, cells were washed once in 1x PBS and infected with CHIKV WT or nsP3-mCherry tagged CHIKV at MOI = 1 in RPMI 1640 media containing 2% FBS and incubated for 1 hour with occasional rocking. Virus-containing media was removed and replaced with fresh RPMI 1640 media containing 2% FBS. After 24 hours of infection, the cells were washed three times with pre-warmed PBS and fixed with 4% paraformaldehyde (PFA, Tousimis) for 15 minutes. After three times PBS washing, the cells were then permeabilized with 0.5% Triton X-100 (Sigma) in PBS for 15 minutes and blocked using 5% bovine serum albumin (BSA) (Sigma) in PBS for 1 hour. Subsequently, the samples were stained with anti-dsRNA (1:200, SCICONS) and anti-nsP1 (1:200, a gift from Andres Merits) at room temperature for 1 hour, and then washed three times with PBS, followed by staining with the secondary antibody anti-mouse IgG Alexa 594 (1:200, Life Technologies) and anti-rabbit IgG Alexa 488 (1:200, Life Technologies) at room temperature for 1 hour. The samples were then washed with PBS three times for subsequent imaging.

### 12. Expansion microscopy (ExM) on nanostructure array

The ExM sample preparation followed a modified version of the protocol as described by Tillberg et al.^75^ and Nakamoto et al.^76^ To make the cell membrane wrap tightly around the nanostructure, the surface of the nanochip was first cleaned with high-power Air-plasma (Harrick Plasma) treatment for 1 hour and then coated with 50 μg/ml poly-L-lysine (PLL, Sigma) for 20 minutes, 0.5% glutaraldehyde (Sigma) for 30 minutes and 0.02 mg/ml gelatin (Sigma) labeled by Atto647N-NHS ester (Sigma) for 40 minutes. For poor gelatin coating around the nanostructure, as shown in **Fig. 5k, l**, plasma cleaning is not necessary. U2OS cells were then cultured on the nanochips. After CHIKV infection and immunostaining, cells were incubated overnight at 4 degrees in a 1: 100 dilution of Acryloy-X (Life Technologies) in PBS. Subsequently, cells were incubated in the gelation solution (monomer solution (8.6% (w/w) Sodium acrylate (Sigma), 2.5% (w/w) Acrylamide (Sigma), 0.15% (w/w) N, N′-Methylenebis (acrylamide) (Sigma)) with 0.2% Tetramethylethylenediamine (TEMED, Sigma), 0.01% 4-Hydroxy-TEMPO (Sigma) and 0.2% Ammonium persulfate (APS, Sigma) in four degrees for overnight. After that, the nanochip with the hydrogel was incubated in digestion buffer (50 mM Tris HCl, 1 mM EDTA, 0.5% Triton X-100, 1 M NaCl) with a 1: 100 diluted proteinase K (Life Technologies). After 2 hours of incubation at room temperature, the hydrogel was detached from the nanochip. Hydrogel was then incubated overnight in MiliQ water at 4 degrees. Before imaging, the hydrogel was carefully moved to a PLL-coated glass bottom dish with a few drops of MiliQ water to keep the humidity.

### 13. Microscopy imaging

Imaging of transfection results in live cells, the time series of nsP1 dynamics in live cells, the bacterial expressed nsP1 monomer (eGFP-nsP1-(b)) distributed on SLB nanobar structures, and the related FRAP tests were performed with laser scanning confocal microscopy (Zeiss LSM 800 with Airyscan) at 100x/1.4 oil objective. For live cell imaging, the cells were maintained in live cell imaging solution (Life Technologies) at 37 ℃ in an on-stage incubator. For *in-vitro* SLB imaging, the free chips were attached to the PDMS chamber. EGFP and Alexa Fluor™ 488-labeled nsP1 were excited at 488 nm and detected at 490-570 nm. Texas-Red-PE containing lipid bilayer was excited at 561 nm and detected at 570-645 nm. CellMask^TM^ Deep Red was excited at 633 nm and detected at 645-700 nm. Each image had a resolution of 512×512 pixels, with a pixel size of 124 nm and a bit depth of 16. The scanning speed was 20s/image with an averaging mode of 4 times/line. For time series imaging, the scanning speed was 1s/image without averaging mode. For FRAP imaging, a circle area with one nanobar was selected and bleached with laser. FRAP videos were collected with 1s/image without averaging mode.

The STED super-resolution nsP1 imaging in cells was performed with a STEDYCON scanner (Abberior Instruments) with 488, 561, and 640 nm excitation lasers together with a 775 nm STED laser (all pulsed). The STEDYCON was mounted to a Nikon Ti2 inverted microscope at 100x/1.45 oil objective. Before imaging, the cells were stained with nsP1-specific antibody, followed by STAR RED (Abberior) secondary antibody, and finally mounted with MOUNT media (Abberior).

Imaging of the FRAP test of the nsP1 and its mutations around nanobars in live cells, the lipid charge effect, the IDR (a.a. 477-516) effect on bacterial expressed nsP1 curvature sensing, the mammalian expressed nsP1 and its MA loops mutations’ distributed on SLB - nanobar structures, and the CHIKV infection in cells around nanostructures with or without ExM were performed with a spinning disc confocal microscope (SDC) that building around a Nikon Ti2 inverted microscope equipped with a Yokogawa CSU-W1 confocal spinning head at 100x/1.4 oil objective. EGFP was excited at 488 nm and detected at 490-570 nm. CellMask Deep Red was excited at 639 nm and detected at 672-712 nm. Each image had a resolution of 1152×1152 pixels, with a pixel size of 65 nm. For FRAP imaging, a circle area with one nanobar was selected and bleached with laser. FRAP videos were collected with 2s/image.

### 14. Image analysis and quantification

To quantify the protein membrane curvature response in live cells, the end-to-center ratio was calculated using ImageJ and MATLAB code as described in previously reported work.^28^ In brief, the background of each individual nanobar image was first corrected by subtracting the local background. with the rolling ball algorithm in ImageJ (radius = 3 pixels for 250 nm diameter nanobars). Next, the individual nanobar position was located using a square mask with the position of the end area and center area. Nanobars with signals from random protein aggregation or with undetectable protein signals were excluded for later processing. To minimize human bias, all the parameters for the calculation and the sorting criteria for data validation were kept consistent. The average intensity values of each square mask were used to generate the average images. Intensity values of pixels within each end and center area were extracted to get the end-to-center ratio. The kymograph and the migration of nsP1 protein clusters were manually tracked using ImageJ, and the protein migratory behavior was characterized using the method as a previous work described.^77^ To generate the FRAP recovery curve, the FRAP area was selected using ImageJ, and the intensity was plotted each time with bleaching correction. Statistical analysis was performed by PRISM 9 (GraphPad). The membrane binding abilities between nsP1-WT and nsP1-W258A were compared by drawing a line profile from the background to the cytosol of the cell and plotting the intensity along this line using imageJ. The intensity was then normalized from 0 to 100, and the mean normalized intensity at each position was plotted.

For *in-vitro* SLB assay curvature sensing quantification, we first located the square mask for individual nanobars. After that, the background of each nanobar image was subtracted using the meshgrid function in Matlab based on the four ROIs at the corners of the image to adjust the uneven surface problem. The average images and end-to-center ratios were then collected as the live cell assay described. For gradient nanobar quantification, the intensities of background-corrected individual nanobar images with different sizes were first normalized to the percentage of the largest-sized nanobar-center intensity to decrease the batch variation. Protein density was measured by the ratio of normalized protein intensity to normalized lipid intensity. The nanobar end areas were adjusted according to the dimension of the nanobar to minimize the covering of the nearby non-curved center of the nanobar to get the normalized nanobar end density.

The density of dsRNA within the inner ring of the nanoring was quantified by measuring the dsRNA signal per unit membrane area, marked by PM staining. The nearby flat area was located 40 pixels to the right of each inner ring. For different-sized inner rings, the dsRNA intensity per unit area was quantified by measuring the integrated dsRNA signal within the inner ring and dividing it by the corresponding inner ring area. Similarly, the dsRNA intensity per unit perimeter was calculated by dividing the dsRNA signal by the respective rim perimeter. Statistical analysis was performed by PRISM 9 (GraphPad).

### 15. Molecular dynamics simulations of nsP1 protein-lipid membrane interaction

#### Simulation of nsP1 monomer-membrane interaction

The interaction between a single copy of nsP1 protein with a planar membrane was modeled using all-atom and coarse-grained molecular dynamic MD simulation. For all-atom simulation, the atomic coordinates of the nsP1 monomer were taken from the dodecamer structure from the Protein Data Bank (code 7DOP).^45^ The missing residues (366 to 375, 415 to 418, and 452 to 458) in the nsP1 monomer structure were modeled using DaReUs-Loop Webserver.^78^ Missing residues 31 to 35 could not be modeled by the webserver so we omitted the first 35 residues from the model as they form an independent segment that is distal from the MA loops and also not involved in the inter-monomer interface. The 3D model was then placed in a simulation box with TIP3P water molecules and neutralizing chloride ions under 3D periodic boundary conditions. MD simulation was next performed using GROMACS version 2018.2 with the all-atom CHARMM27 forcefield.^79–81^ After energy minimization and equilibration with positional restraints on non-H atoms, unrestrained simulation was performed for 5 ns with a 2 fs time-step to equilibrate the nsP1 monomer in water. The temperature was maintained at 303 K and pressure was maintained at 1 bar. Electrostatic interactions were computed using Particle Mesh Ewald method with a cut-off distance of 1.2 nm. Van der Waals (vdW) interactions were computed using a cut-off method with forces smoothly switched to zero between 1.0 and 1.2 nm. The equilibrated nsP1 monomer structure was then placed on top of an equilibrated planar membrane composed of 80% POPC and 20% POPS lipids. The lipid membrane was built using Bilayer Membrane Builder in CHARMMM-GUI webserver^82^ consisting of 230 lipids per leaflet. The system was solvated with TIP3P water molecules and with 3D periodic boundary conditions imposed. Sodium and chloride ions were added to electrically neutralize the system. CHARMM36m force field was used to describe inter-atomic interactions. Energy minimization was then performed to avoid any clashes between the molecules followed by equilibration of the system for 2 ns at 310 K and 1 bar. Three independent simulations were carried out for 20 ns by assigning different sets of initial velocities to the atoms (**Supplementary Fig. 23**).

To analyze the interactions between nsP1 monomer and lipid membrane at longer time scales, coarse-grained simulations were performed. For coarse-grained simulations, the lipid membrane along with water and ions were coarse-grained with the help of martinize script following MARTINI model of coarse-graining.^83^ The nsP1 protein was coarse-grained separately and then placed over the lipid membrane. After setting up the initial configuration, the system was energy minimized followed by equilibration for 5 ns. Production run was carried out for 300 ns with temperature maintained at 310 K and pressure maintained at 1 bar. Velocity-rescale method was used for temperature coupling, while pressure coupling was performed using Berendsen method and Parrinello-Rahman method during equilibration and production run, respectively. The simulations were repeated three times with different initial velocities assigned to coarse-grained beads similar to all-atom simulations discussed above.

#### Simulation of nsP1 dodecamer-membrane interaction

Further, the interaction between the nsP1 dodecamer and planar membrane was analyzed using coarse-grained MD simulation for computational efficiency. Firstly, the complete nsP1 monomer model was fitted to each of the 12 subunits in the dodecamer to obtain a complete all-atom model. Next, a coarse-grained nsP1 dodecamer structure was obtained based on the MARTINI coarse-grained model using *martinize* script version 2.6.3.^83^ The coarse-grained nsP1 dodecamer was then placed on top of a planar lipid membrane (2000 lipids) with POPC: POPS lipids (80%:20%) in the upper leaflet using CHARMM-GUI interface. The lipid membrane was modeled using MARTINI 3.0 forcefield.^84^ After energy minimization, equilibration for 5 ns was conducted at 310 K and 1 bar, followed by a production run for 1400 ns at the same temperature and pressure. To quantify changes to the membrane curvature upon nsP1 ring binding, the *g_lomepro* tool downloaded from http://www3.mpibpc.mpg.de/groups/de_groot/g_lomepro.html was used to obtain the local mean membrane curvature.^85^ Simulation snapshots from 1360 to 1400 ns (40 frames), during which the nsP1 ring was stably bound to the membrane, were used for the analysis. A band-pass filter (*q*low = 0.3 nm^−1^ and *q*high = 0.7 nm^−1^) was applied to the computed curvature modes so as to capture the strongly curved region of the membrane. Finally, simulations were set up to analyze the binding affinity of nsP1 dodecamer towards SLBs with and without curvature. A flat membrane and a curved one with a radius of curvature of 100 nm were set up following the protocol by Liu et. al..^86^ In brief, the lipid head beads in the inner leaflet were fixed with the help of a restraining force constant of 1000 kJ mol^-1^ nm^-2^. The simulations were conducted using coarse-grained MD simulation technique with MARTINI 3.0 forcefield and similar conditions as mentioned in previous coarse-grained planar membrane simulations. Finally, it’s the total interaction energy between nsP1 dodecamer and with both planar and curved membranes. This is one of the commonly used methods to quantify the binding strength between different molecules of interest.^87–90^ In addition, the insertion depths of MA loops in the planar and curved membranes were quantified with the help of a TCL script.

