## supplementary information for "Saddle Curvature Association of nsP1 Facilitates the Replication Complex Assembly of Chikungunya Virus in Cells"

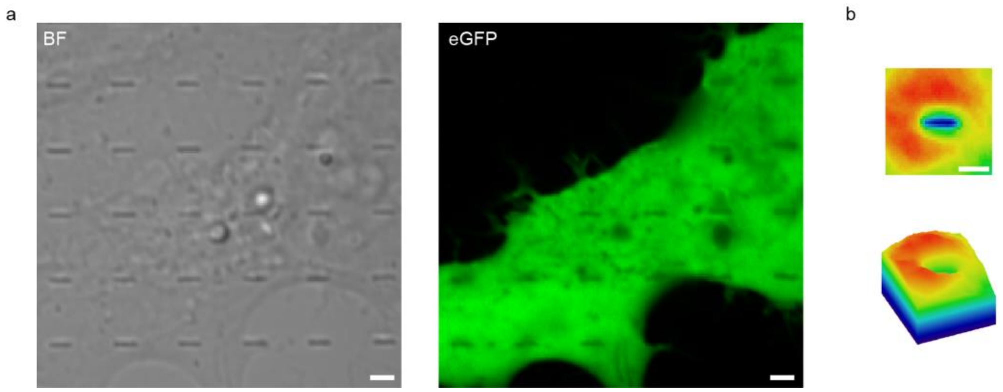


**Suppl. Figure S1. GFP distribution around nanobars in U2OS cells.** (a) Representative image of a cell transfected with GFP. Scale bars: 2 μm. (b) Average image and corresponding 3D surface plot of the GFP distribution around nanobars. n = 134 bars.

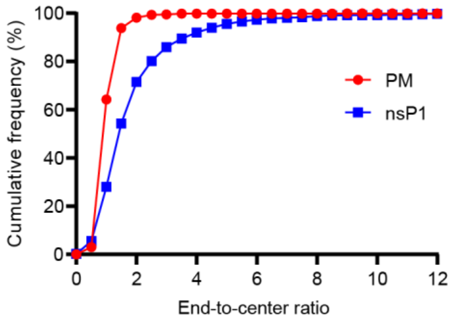


**Suppl. Fig. S2. Accumulated frequency distribution of membrane and eGFP-nsP1 end-to-center ratio.** n = 995 bars for PM and n =1005 bars for eGFP-nsP1, with 3 independent experiments.

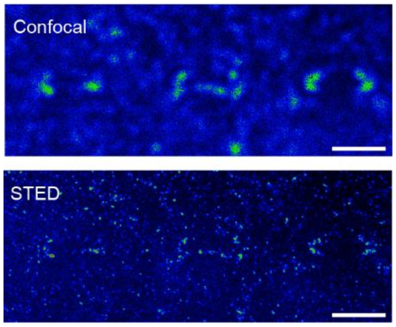


**Suppl. Fig. S3. The accumulation of nsP1 at positively curved nanobar ends imaged by either confocal (top) or STED (bottom) microscopy.** Scale bars: 2 µm.

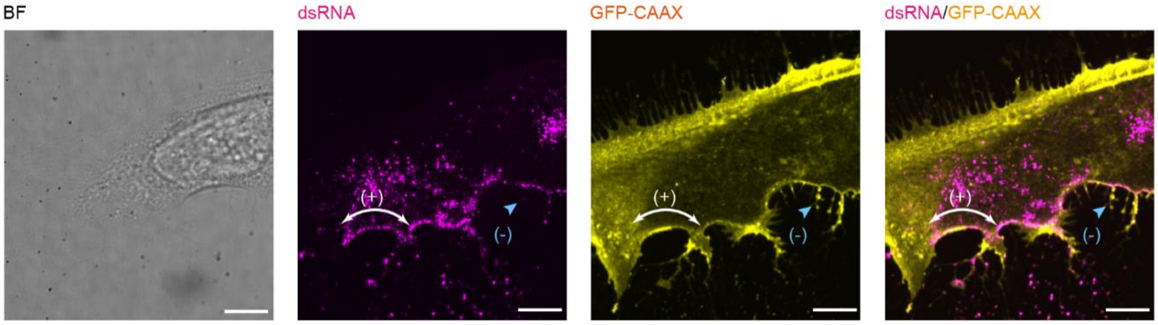


**Suppl. Fig. S4. Localization of the replication sites in a CHIKV-infected cell.** A representative image of a cell cultured on a flat surface and infected with CHIKV. The cell was transfected by GFP-CAAX (yellow) to mark the cell membrane and stained with anti-double-stranded RNA (dsRNA, magenta) to indicate viral replication sites. The result showed that CHIKV replication preferred to accumulate at the positively curved membrane of the host cell (white arrow), compared to the protruded negatively curved membranes (blue arrow). n = 13 cells. Scale bars: 10 μm.

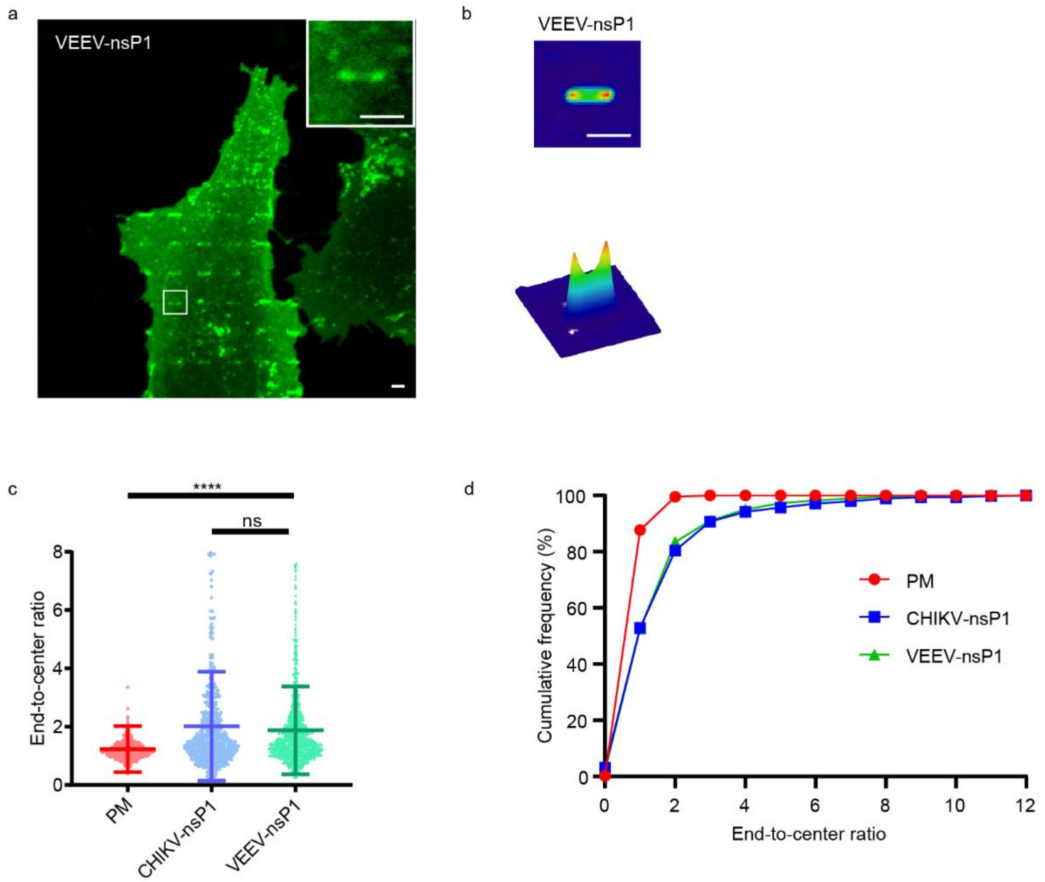


**Suppl. Figure S5. VEEV-nsP1 curvature sensing ability in U2OS cells.** (a) Representative fluorescence image of a cell transfected with VEEV-nsP1. Scale bar: 2 μm. (b) Average image and corresponding 3D surface plot of the VEEV-nsP1 distribution around nanobars. n = 597 bars for VEEV-nsP1. (c) The end-to-center ratio of the PM, CHIKV-nsP1, and VEEV-nsP1 around nanobar. Data are the mean ± SD. P value calculated using unpaired t-test with Welch's correction: **** represents $p < 0.0001$. n = 439, 489, 1097 bars respectively, with 3 independent experiments. (d) Accumulated frequency distribution of PM, CHIKV-nsP1, and VEEV-nsP1 end-to-center ratio. n = 439, 489, 1097 bars respectively, with 3 independent experiments.

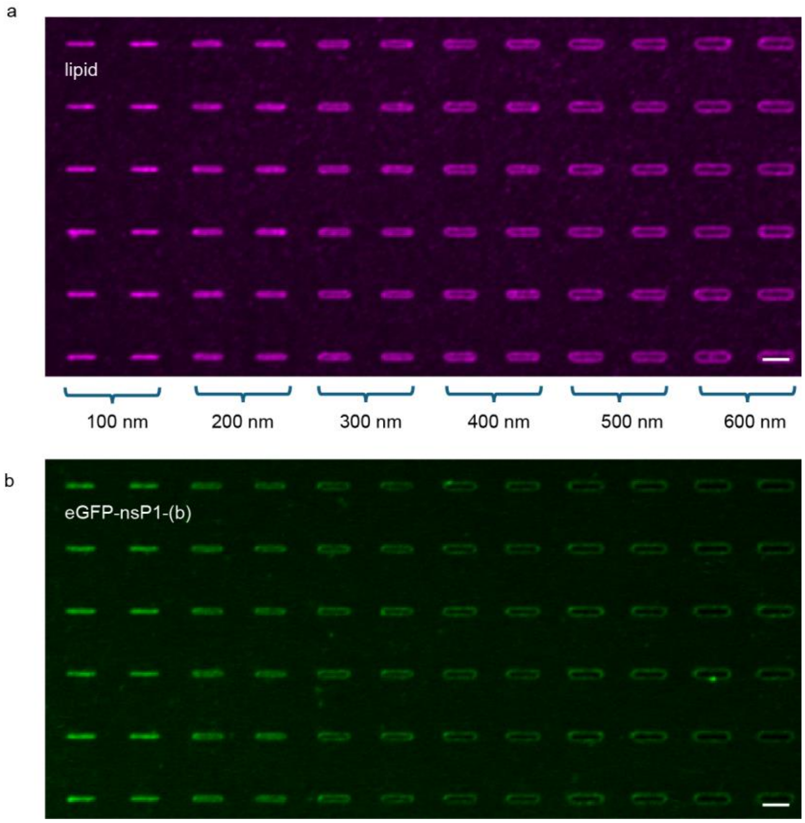


**Suppl. Figure S6. eGFP-nsP1-(b) curvature sensing ability *in vitro*.** (a) Representative image of SLB (89.5% egg PC + 10% brain PS + 0.5% Texas-Red-PE) on nanobar array with increasing width from 100 nm to 600 nm. (b) Representative image of the eGFP-nsP1-(b) bonded to SLB on nanobar array with increasing width from 100 nm to 600 nm. Scale bars: 2 μm.

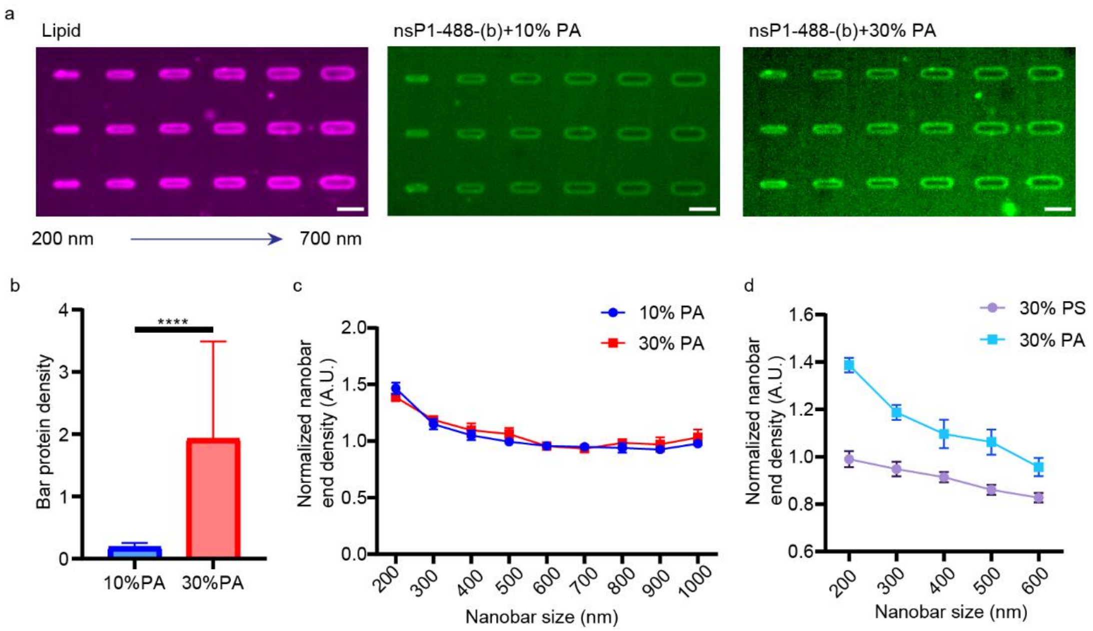


**Suppl. Figure S7. The nsP1 curvature sensing ability on PA-based SLB.** (a) Representative images of 1 μM Alexa Fluor 488 labeled nsP1 (green) on 10% PA + 89.5% POPC + 0.5% Texas-Red-PE SLB or 30% POPS + 69.5% POPC + 0.5% Texas-Red-PE SLB (magenta) on gradient nanobars with width from 200 to 1000 nm. Scale bars: 2 μm. (b) Relative protein density of nsP1 for different lipid compositions over 80 bars. Data are the mean ± SD. P value calculated using unpaired t-test with Welch's correction: **** represents $p < 0.0001$, with 3 independent experiments. (c) Normalized nanobar-end density of nsP1 based on the corresponding lipid bilayer intensity on 10% PA and 30% PA SLB over 80 bars. Data are the mean ± SEM, with 3 independent experiments. (d) Comparison of nsP1 curvature sensing ability on 30% PA and 30% PS. Data are the mean ± SEM, with 3 independent experiments.

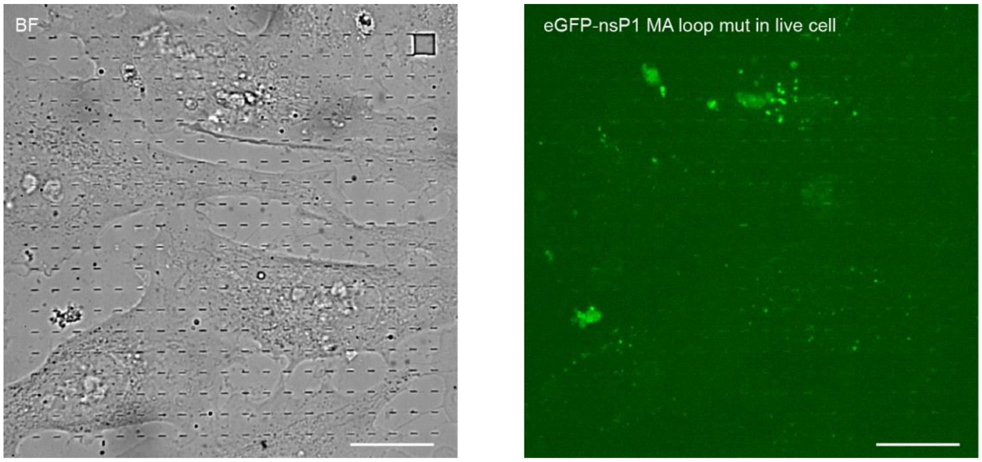


**Suppl. Figure S8. The expression of the nsP1 MA loops mutant in U2OS cells.** Representative brightfield and fluorescence images of the U2OS cells transfected with nsP1 MA loops mutant. The result showed that the nsP1 MA loops mutant is hardly expressed in the U2OS cells. Scale bars: 20 μm.

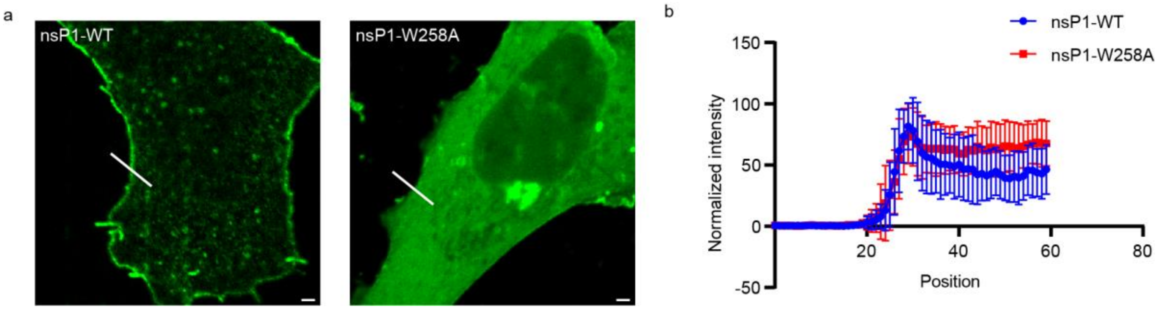


**Suppl. Figure S9. Membrane binding ability of nsP1-WT *vs* nsP1-W258A.** (a) Representative images of nsP1-WT and nsP1-W258A expressed in U2OS cells. Scale bars: 2 μm. (b) The normalized intensity of the line profile (an example is the white line in panel a) from the background to the cytosol over 10 cells. Data are the mean ± SD.

a

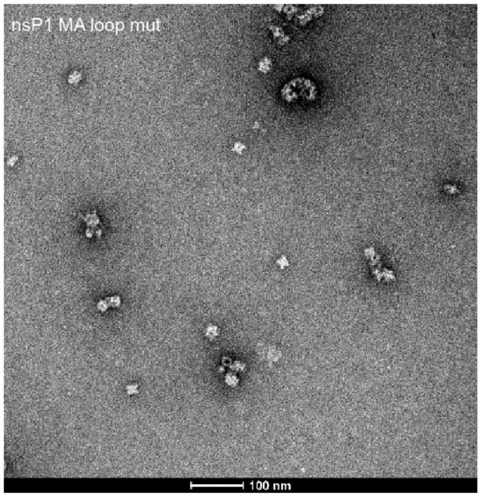


b

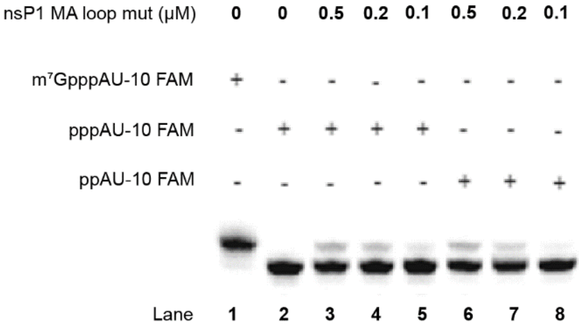


**Suppl. Figure S10. Characterization of purified protein nsP1 MA loop mutant.** (a) The negative-staining EM image of nsP1 MA loop mutant. Scale bars: 100 nm. (b) Capping activity of nsP1 MA loop mutant.

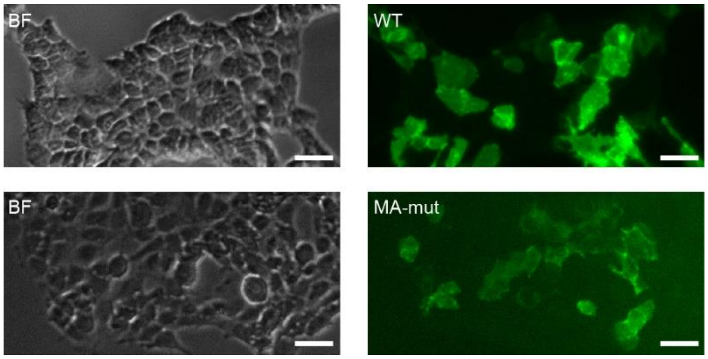


**Suppl. Figure S11. The expression of P1234 and P1234-MA-mut in HEK293-FT cells.** The expression of P1234 and P1234-MA-mut in HEK293-FT cells was successfully confirmed by nsP1 immunostaining. Scale bars: 40 μm

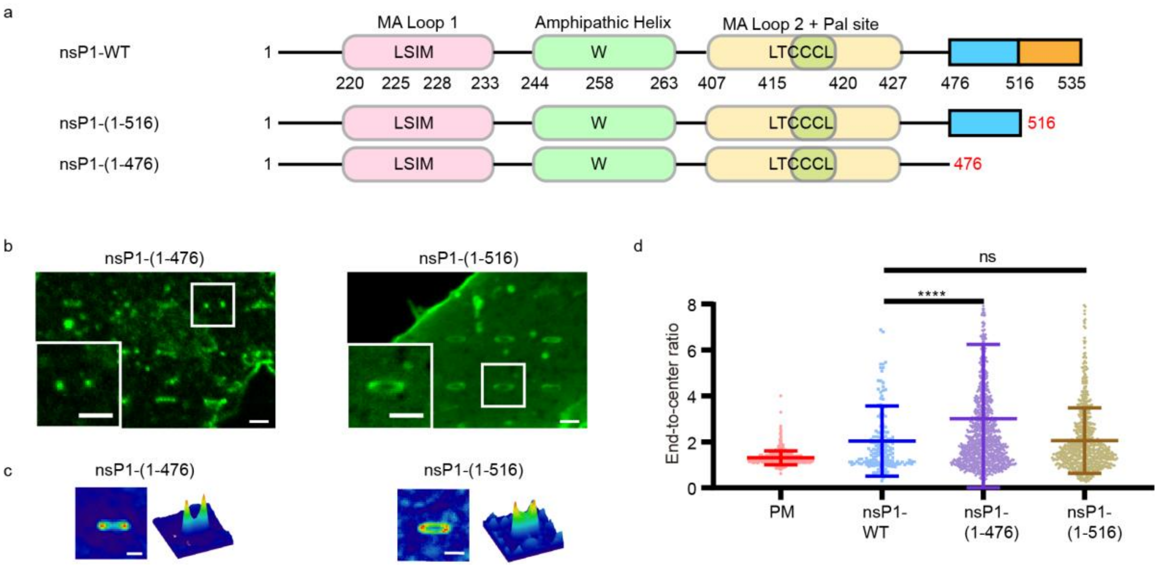


**Suppl. Figure S12. CHIKV nsP1 curvature sensing ability with the c-terminal disorder domain.** (a) The CHIKV nsP1 truncations at the c-terminal disordered regions which were used in this study. (b) Representative images of cells cultured on nanobars expressing eGFP tagged nsP1-(1-476) (left) or nsP1-(1-516) (right). (c) Average images and surface plots of U2OS cells cultured on nanobars expressing eGFP tagged nsP1-(1-476) and nsP1-(1-516). n = 697 bars for nsP1-(1-476), and n = 451 bars for nsP1-(1-516). (d) The end-to-center ratio of the PM, nsP1-WT, nsP1-(1-476) and nsP1-(1-516) on nanobars. Data are the mean ± SD. P value calculated using unpaired t-test with Welch's correction: **** represents $p < 0.0001$. n = 539, 170, 1129, 743 bars respectively, with 3 independent experiments. Scale bars: 2 μm.

Positive Charge    Negative Charge

MDPVYVDIDADSAFLKALQRAYPMFEVEPRQVTPNDHANARAFSHLAIKLIEQEIDPDSTILDIGSAPARRMMSDRKYHCVCPM
RSAEDPERLANYARKLASAAGKVLDRNISGKIGDLQAVMAVPDKETPTFCLHTDVSCRQRADVAIYQDVYAVHAPTSLYHQAIK
GVRVAYWVGFDTTPFMYNAMAGAYPSYSTNWADEQVLKAKNIGLCSTDLTEGRRGKLSIMRGKKLKPCDRVLFSVGSTLYPE
SRKLLKSWHLPSVFHLKGKLSFTCRCDTVVSCEGYVVKRITMSPGLYGKTTGYAVTHHADGFLMCKTTDTVDGERVSFSVCT
YVPATICDQMTGILATEVTPEDAQKLLVGLNQRIVVNGRTQRNMNTMKNYLLPVVAQAFSKWAKECRKDMEDEKLLGVRERTL
TCCCLWAFKKQKTHTVYKRPDTQSIQKVQAEFDSFVVPSLWSSGLSIPLRTRIKWLLSKVPKTDLIPYSGDAREARDAEKEAEE
477
EREAELTREALPSLQAAQEDVQVEIDVEQLEDRAGA
516

**Suppl. Figure S13. CHIKV nsP1 full-length sequence.** The sequence between 477 to 516 is highlighted in orange underline, including the blue color highlighted positively charged amino acids and the pink color highlighted negatively charged amino acids.

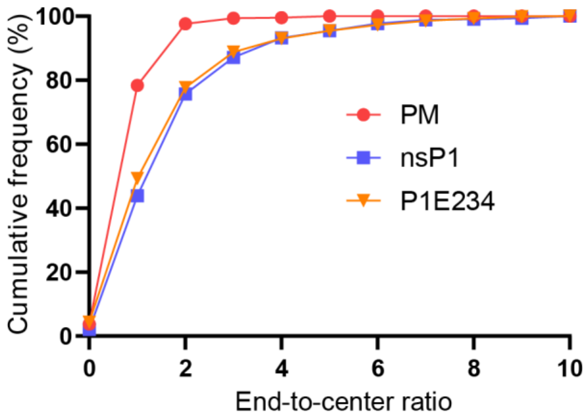


**Suppl. Figure S14. Accumulated frequency distribution of P1E234 end-to-center ratio.** The accumulated frequency distribution of the P1E234 end-to-center ratio shows no significant difference compared to nsP1-WT. n = 666, 626, 1491 bars respectively, with 3 independent experiments.

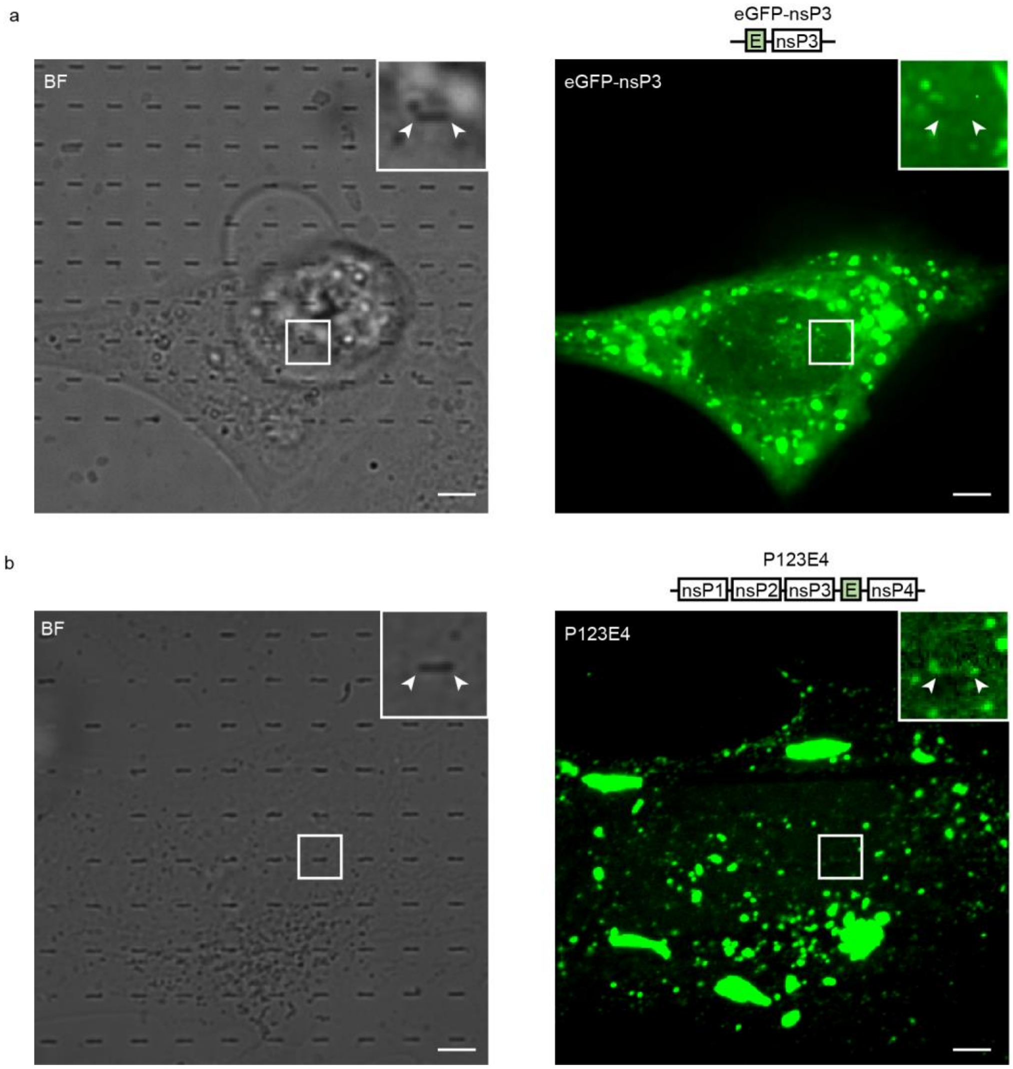


**Suppl. Figure S15. CHIKV nsP3 binds to the membrane via the association with nsP1.** (a) Representative images of a cell transfected with eGFP-nsP3 (green) on the nanobars in brightfield and fluorescence. (b) Representative images of U2OS cells transfected with P123E4-WT (green) on the nanobars in brightfield and fluorescence. Scale bars: 5 µm.

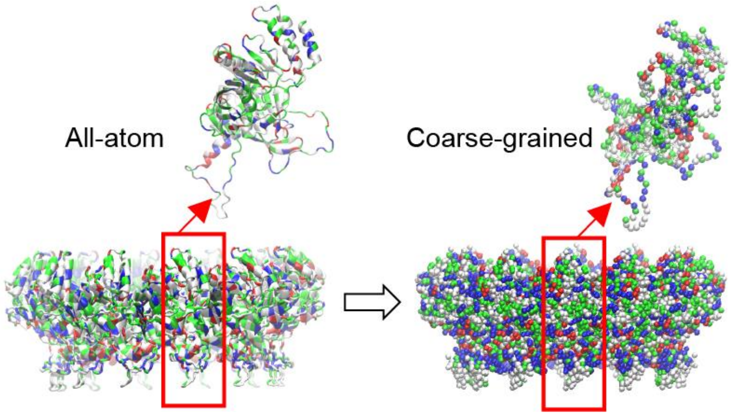


**Suppl. Figure S16. Coarse-grained nsP1 dodecamer from the all-atom structure.** The red box area represents a single copy of nsP1 from the dodecamer.

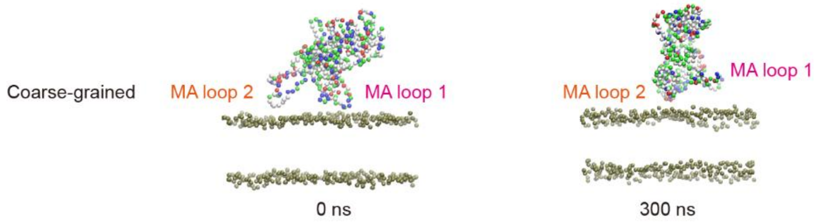


**Suppl. Figure S17. Coarse-grained single nsP1 on the planar membrane.** Snapshot of a coarse-grained simulation showing the interaction of a single copy of nsP1 protein with a planar membrane at initial (left) and equilibrated (right) states.

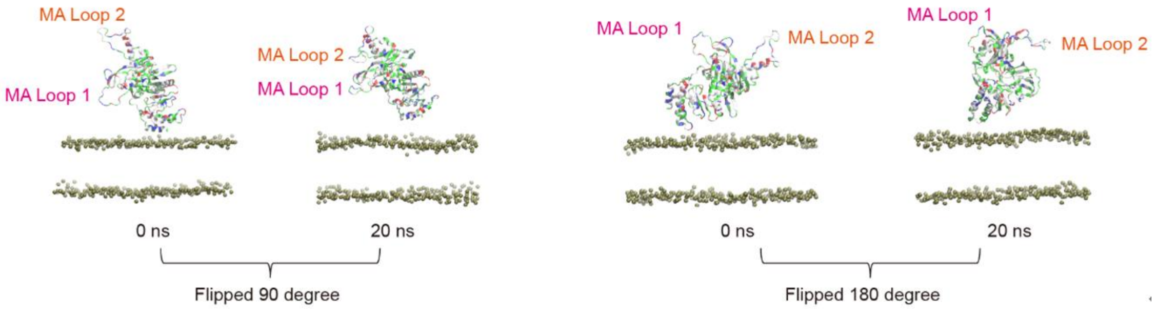


**Suppl. Figure S18. Flipped single nsP1 on the planar membrane.** Snapshot of a coarse-grained simulation showing the interaction of a single copy of nsP1 which flipped 90 degrees and 180 degrees with a planar membrane at initial (left) and equilibrated (right) states.

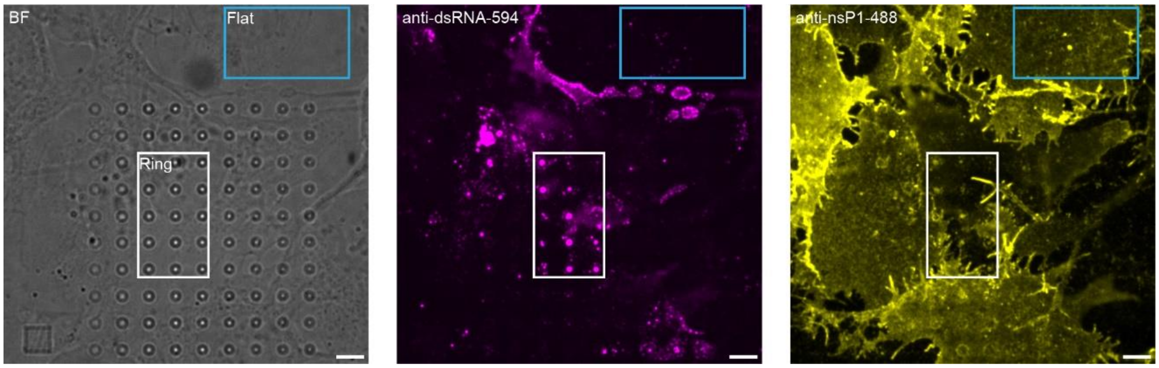


**Suppl. Figure S19. U2OS cells infected with WT CHIKV on nanoring arrays.** Representative brightfield and fluorescence images of CHIKV infected cells on nanoring arrays. Cells were stained for dsRNA (magenta) and nsP1 (yellow). Zoomed-in images of the brightfield and dsRNA within the white boxes are shown in **Fig. 5c.** The nsP1 staining indicated that nsP1 was also present at the replication sites, suggesting its involvement in the CHIKV replication. Scale bars: 5 µm.

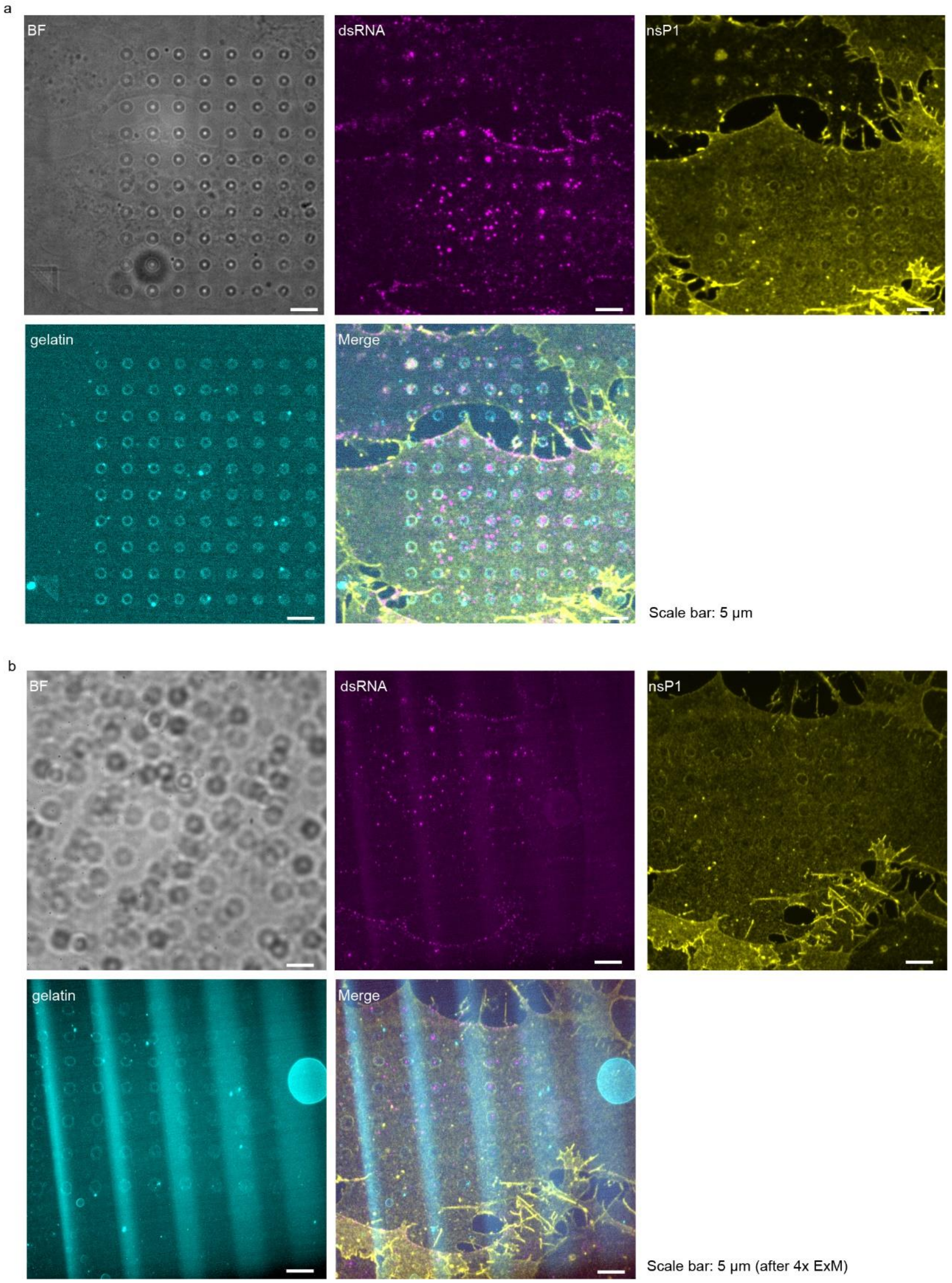


**Suppl. Figure S20. Pre-and post-ExM fluorescent imaging.** (a) Representative fluorescence images of a CHIKV-infected cell cultured on nanoring arrays with a gelatin-coated surface (cyan). The cells were stained for dsRNA (magenta) and nsP1 (yellow). (b) The same cell after ~4x ExM.

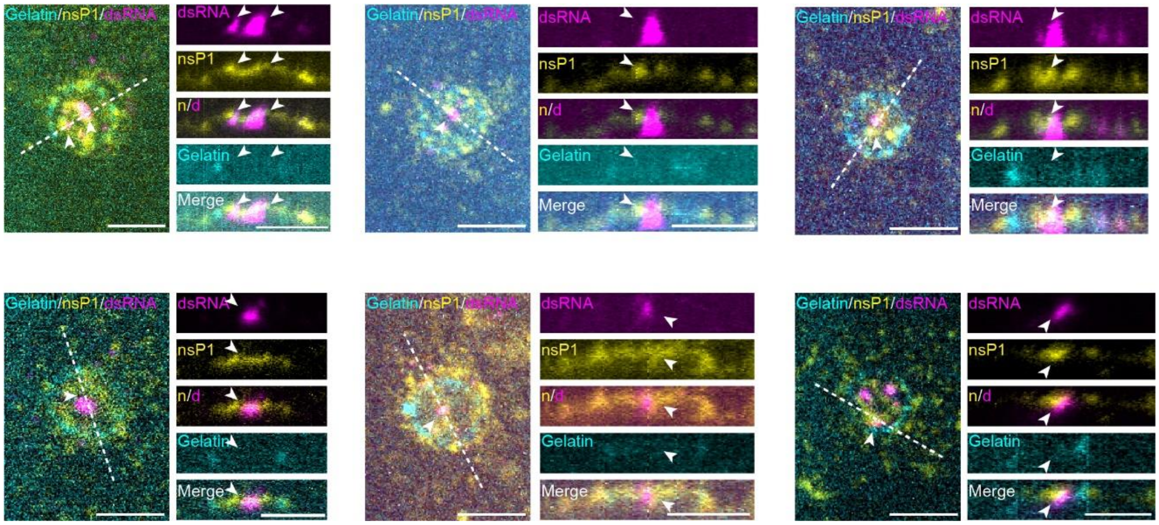


**Suppl. Figure S21. Zoomed-in examples of the post-ExM images with cells cultured on nanorings.** Left: the x-y image of the nanoring coated by gelatin (cyan). Cells were stained for dsRNA (magenta) and nsP1 (yellow). Right: the resliced image along the white line displayed the view across the z-axis. Scale bars: 8 µm.

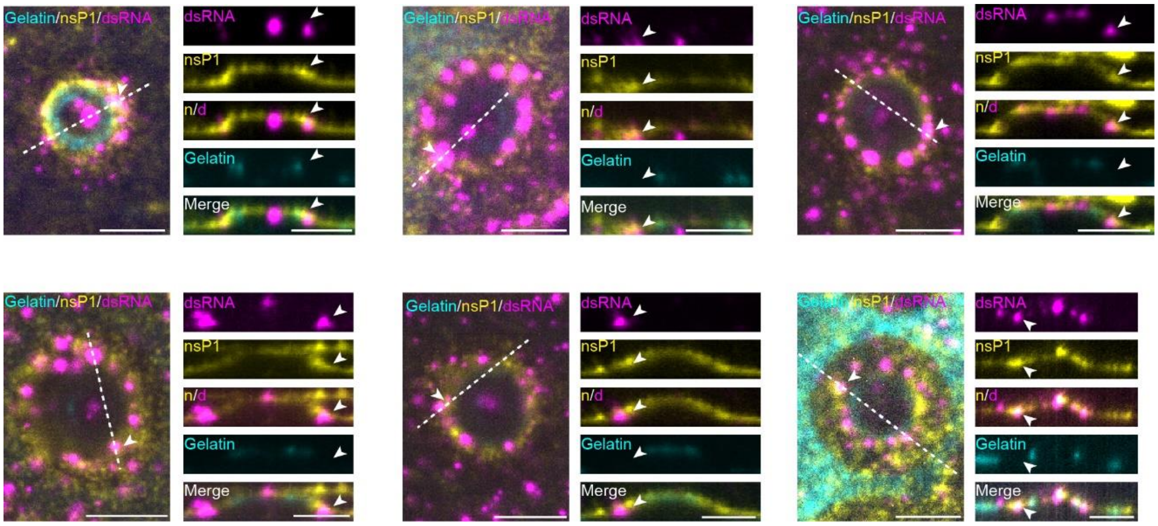


**Suppl. Figure S22. Zoomed-in examples of the post-ExM images showing cells cultured on nanorings with incomplete PM adhesion.** Left: the x-y image of the nanoring coated by gelatin (cyan). Cells were stained for dsRNA (magenta) and nsP1 (yellow). Right: the resliced image along the white line displayed the view across the z-axis. Scale bars: 8 μm.

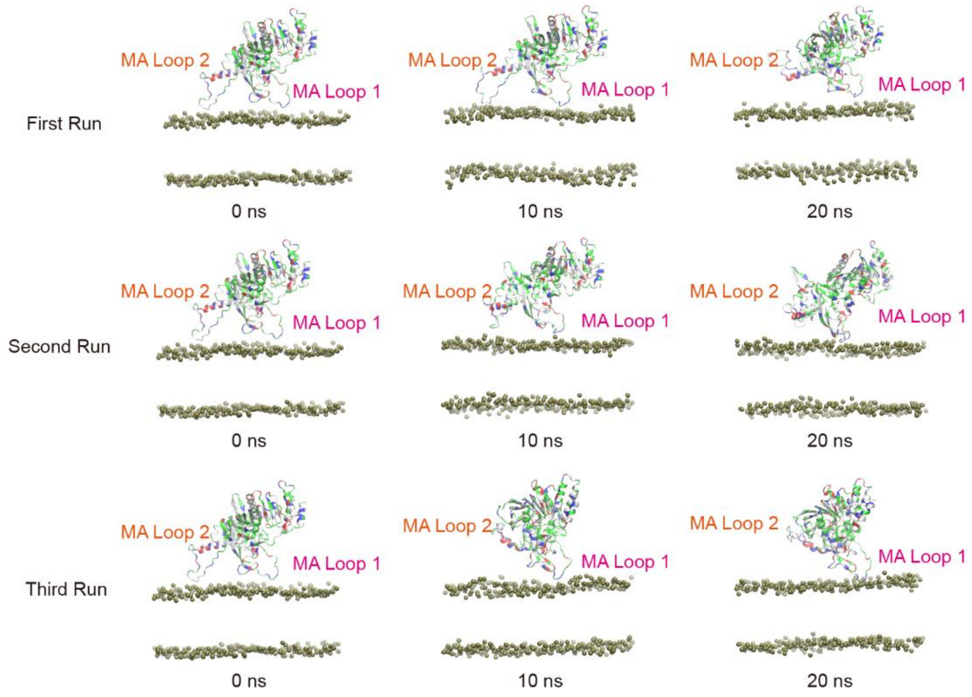


**Suppl. Figure S23. Three independent simulations of nsP1 protein-lipid membrane interaction.** The results of three independent all-atom MD simulations show the interaction of a single copy of nsP1 protein with a planar membrane.

### List of Supplementary Movies:

**Movie 1**. **EGFP-nsP1 distribution around nanobars.** Time-lapses of eGFP-nsP1 distribution around two adjacent nanobars in six minutes. Scale bars: 2 μm.

**Movie 2**. **FRAP test of the PM and eGFP-nsP1 around nanobar.** After photobleaching, the CellMask Deep Red stained PM (magenta) around the nanobar was recovered, while the overexpressed eGFP-nsP1 (green) showed no obvious recovery during 120 s. Scale bars: 2 μm.

**Movie 3. FRAP test of SLB and purified nsP1 around nanobar.** After photobleaching, both SLB (10% Brain PS + 89.5% Egg PC + 0.5% Texas Red DHPE) and bacterial expressed nsP1 around the nanobar were recovered during 90 s. Scale bars: 2 μm.

**Movie 4**. **FRAP tests of the overexpressed eGFP-nsP1, eGFP-nsP1-W258A, and eGFP-nsP1-CCC_3A around nanobars.** EGFP-nsP1-WT and eGFP-nsP1-W258A around nanobars showed no detectable recovery after photobleaching. In comparison, eGFP-nsP1-CCC_3A around nanobar was recovered during 120 s. Scale bars: 2 μm.

**Movie 5**. **Coarse-grained simulation of nsP1 binds to the membrane.** The binding of nsP1 dodecamer generated local positive membrane curvature on the initially flat membrane surface.

**Movie 6. The z-stack ExM image of a CHIKV-infected cell on the nanoring array.** The surface of the nanoring was coated by fluorescence-labeled gelatin (cyan). The cells were stained with anti-nsP1 (yellow) and anti-dsRNA (magenta). The z-stack was from the cell bottom to the cell top. Scale bar: 20 μm.
